# A genetically encoded Ca^2+^ indicator based on circularly permutated sea anemone red fluorescent protein

**DOI:** 10.1101/213082

**Authors:** Yi Shen, Hod Dana, Ahmed S. Abdelfattah, Ronak Patel, Jamien Shea, Rosana S. Molina, Bijal Rawal, Vladimir Rancic, Yu-Fen Chang, Lanshi Wu, Yingche Chen, Yong Qian, Matthew D. Wiens, Nathan Hambleton, Klaus Ballanyi, Thomas E. Hughes, Mikhail Drobizhev, Douglas S. Kim, Minoru Koyama, Eric R. Schreiter, Robert E. Campbell

**Affiliations:** Department of Chemistry, University of Alberta, Edmonton, Alberta T6G 2G2, Canada.; Janelia Research Campus, Howard Hughes Medical Institute, Ashburn, Virginia 20147, USA.; Department of Cell Biology and Neuroscience, Montana State University, Bozeman, Montana 59717, USA.; Department of Physiology, University of Alberta, Edmonton, Alberta T6G 2H7, Canada.; LumiSTAR Biotechnology Incorporation, Nangang Dist., Taipei City 115, Taiwan.

## Abstract

Genetically-encoded calcium ion (Ca^2+^) indicators (GECIs) are indispensable tools for measuring Ca^2+^ dynamics and neuronal activities in vitro and in vivo. Red fluorescent protein (RFP)-based GECIs enable multicolor visualization with blue or cyan-excitable fluorophores and combined use with blue or cyan-excitable optogenetic actuators. Here we report the development, structure, and validation of a new red fluorescent Ca^2+^ indicator, K-GECO1, based on a circularly permutated RFP derived from the sea anemone Entacmaea quadricolor. We characterized the performance of K-GECO1 in cultured HeLa cells, dissociated neurons, stem cell derived cardiomyocytes, organotypic brain slices, zebrafish spinal cord in vivo, and mouse brain in vivo.

## Introduction

Protein engineering efforts have yielded three major lineages of monomeric red fluorescent proteins (RFPs) derived from their naturally oligomeric precursors (**Fig. 1a**). One lineage comes from the *Discosoma sp*. mushroom coral RFP, DsRed, and includes the first monomeric RFP, mRFP1 (Ref. 1), and the mRFP1-derived “mFruit” variants such as mCherry, mCherry2, mOrange, and mApple^2–4^. The second and third lineages stem from the sea anemone *Entacmaea quadricolor* RFPs eqFP578 (Ref. 5) and eqFP611 (Ref. 6), respectively. EqFP578 is the progenitor of the bright monomeric proteins TagRFP, TagRFP-T, mKate, mKate2, and the low cytotoxicity variant FusionRed^57–9^. Engineering of eqFP611 produced mRuby, mRuby2, and mRuby3, a line of RFPs with relatively large stokes-shift and bright red fluorescence^10–12^. Together, these three lineages of monomeric RFPs are commonly used in a variety of fluorescence imaging applications and have served as templates for developing red fluorescent indicators of various biochemical activities^13^.

**Figure 1.**
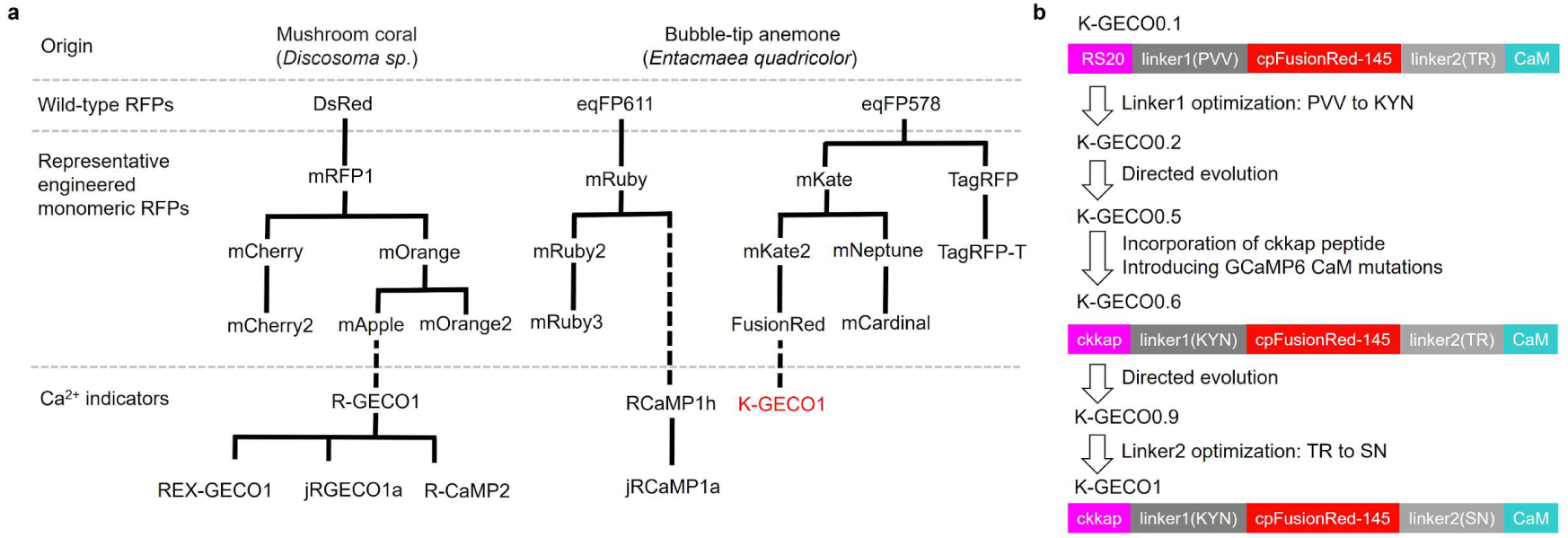
Design and development of K-GECO1. **(a)** Selected RFP and RFP-based Ca^2+^ indicator genealogy. **(b)** Schematic illustration of K-GECO1 design and engineering.

Among the many FP-based indicators of biochemical activity, genetically-encoded calcium ion (Ca^2+^) indicators (GECIs) are particularly versatile tools. Most notably, they enable imaging of neuronal activity in contexts ranging from dissociated neurons *in vitro* to brain activity in behaving animals^14^. Green fluorescent GCaMPs, in particular, have proven extremely useful for imaging Ca^2+^ activities in various neural systems^15–17^. The development of the first single RFP-based Ca^2+^ indicators, the DsRed-derived R-GECO1 (Ref. 18) and eqFP611-derived RCaMP1h^19^, unlocked new opportunities for simultaneous multicolor optical imaging. Further engineering of R-GECO1 produced a number of improved and altered variants, including R-CaMP1.07, R-GECO1.2, CAR-GECO1, O-GECO1, R-CaMP2, and REX-GECO1 (Refs. 20–23). Optimization of R-GECO1 and RCaMP1h for detection of neuronal action potentials produced jRGECO1a, jRCaMP1a, and jRCaMP1b^24^. One limitation of the R-GECO series of GECIs is that they inherited undesirable blue-light activated photoswitching behavior that was also present in the DsRed-derived template (mApple) from which they were engineered^3,19,25,26^. Accordingly, when combining the R-GECO series of Ca^2+^ indicators with optogenetic actuators, extra care must be taken to differentiate true responses from artifacts caused by photoactivation^19,21^. RCaMP variants do not show photoswitching under blue illumination but they are less responsive than R-GECO variants in terms of fluorescence change upon Ca^2+^ binding^19,24^. Like many DsRed-derived RFPs, R-GECO variants have a propensity to accumulate in lysosomes and form brightly fluorescent (but non-functional) puncta during long-term neuronal expression^27–29^. These puncta can complicate image analysis and may compromise long term cell viability.

The drawbacks associated with the DsRed- and eqFP611-derived GECIs motivated us to explore a new RFP template for development of red GECIs. As mentioned above, some DsRed-derived RFPs, such as mOrange and mCherry, have been reported to exhibit relatively dim fluorescence and/or puncta formation, when transgenically expressed in mice brains^30^. In contrast, eqFP578-derived RFPs TagRFP-T and mKate2 have been reported to exhibit bright fluorescence without puncta formation *in vivo^30^*. The eqFP611-derived mRuby has been reported to have the highest cytotoxicity among various RFPs^9^. Based on these literature reports, and reinforced by observations in our own lab, we reasoned that using an eqFP578-derived RFP as a template for the development of a new red GECI could potentially address the limitations of R-GECO, and possibly offer a better performance *in vivo*.

Here we report our efforts to design, engineer, characterize, and validate a new red GECI, K-GECO1, based on the eqFP578 variant FusionRed^9^.

## Results

### Design and engineering of K-GECO1

We initially selected two eqFP578-derived RFPs, mKate2^8^ and its low-cytotoxicity variant FusionRed^9^, as templates for construction of a red Ca^2+^ indicator. Both mKate2 and FusionRed scaffolds were circularly permutated (cp) at residue Ser143 (numbering according to mKate crystal structure^31^, PDB: 3BXB), which is the same permutation site used in GCaMPs and R-GECOs^18,32^. Both cpRFPs were genetically inserted between N-terminal chicken myosin light-chain kinase (MLCK) peptide RS20 and C-terminal Calmodulin (CaM) from R-GECO1. The resulting indicator prototype based on the cpmKate2 scaffold was not fluorescent, in accordance with a previous study of mKate circular permutation^33^, and therefore no further optimization was pursued. In contrast, the cpFusionRed-based design (designated K-GECO0.1) (**Fig. 1b**), was dimly fluorescent when expressed in *Escherichia coli* colonies for 48 h at room temperature. The extracted protein showed 20% fluorescence emission intensity increase upon addition of Ca^2^+. To further improve the function of this prototype indicator, we first performed random mutagenesis of the peptide linker between the RS20 peptide and cpFusionRed (linker1), which is Pro30-Val31-Val32 as in R-GECO1 (numbered as in **Supplementary Fig. 1**). Screening of this targeted mutagenesis library led to identification of the Pro30Lys-Val31Tyr-Val32Asn variant with visible red fluorescence in *E. coli* after overnight incubation. This variant, termed K-GECO0.2, exhibited a 2-fold fluorescence emission intensity increase upon Ca^2^+ binding. K-GECO0.2 was subjected to further directed protein evolution for brightness and larger Ca^2+^-induced fluorescence intensity change. In each round of directed evolution, error prone PCR was used to create a variant library. After visual inspection of the plated library, the brightest fluorescent colonies were picked, cultured, and the protein purified and tested for its Ca^2+^ response. The pool of variants with the largest Ca^2+^dependent fluorescence changes served as templates for the next round of evolution. After three rounds, an improved variant K-GECO0.5 was produced. Initial characterization of K-GECO0.5 indicated a relatively low Ca^2+^ affinity with a *K*_d_ close to 1 μM. In an effort to overcome this limitation, we engineered K-GECO0.6 using an approach similar to the one used by Inoue et al. to develop R-CaMP2 (Ref. 23). Following the strategy of Inoue et al., we incorporated the rat CaM-dependent kinase kinase peptide (ckkap) in place of RS20, and introduced GCaMP6 mutations Asn342Ser, Asp343Tyr, Thr344Arg, and Ser346Thr into the CaM domain^23^. An additional three rounds of directed evolution led to K-GECO0.9. In the final step of engineering, we performed saturation mutagenesis of the linker between cpFusionRed and CaM (linker2). Screening of the library identified a variant with linker2 changed from Thr265-Arg266 into Ser265-Asn266. This final variant was designated as K-GECO1 (Fig. 1b).

### *In vitro* characterization of K-GECO1

The excitation and emission maxima of K-GECO1 are 568 nm and 594 nm, respectively, in the Ca^2+^-unbound state. In the Ca^2+^-bound state, these 2 maxima are slightly blue-shifted to 565 nm and 590 nm (Fig. 2a, Supplementary Table 1. K-GECO1 exhibits a 12-fold fluorescent intensity increase upon Ca^2+^ binding, with extinction coefficient increasing from 19,000 M^−1^ cm^−1^ to 61,000 M^−1^cm^−1^ and quantum yield from 0.12 to 0.45 (Supplementary Table 1). The fluorescence spectra characteristics and Ca^2+^-induced fluorescence change of K-GECO1 are generally very similar to R-GECO1 (Supplementary Table 1). However, K-GECO1 is about 2-fold brighter than R-GECO1 under one-photon excitation. Ca^2+^ titration of purified K-GECO1 reveals that the protein has an apparent *K*_d_ of 165 nM with a Hill coefficient of 1.12 (**Fig. 1b**, **Supplementary Table 1**), similar to R-CaMP2 and other ckkap-based GECIs^23,34^.

**Figure 2.**
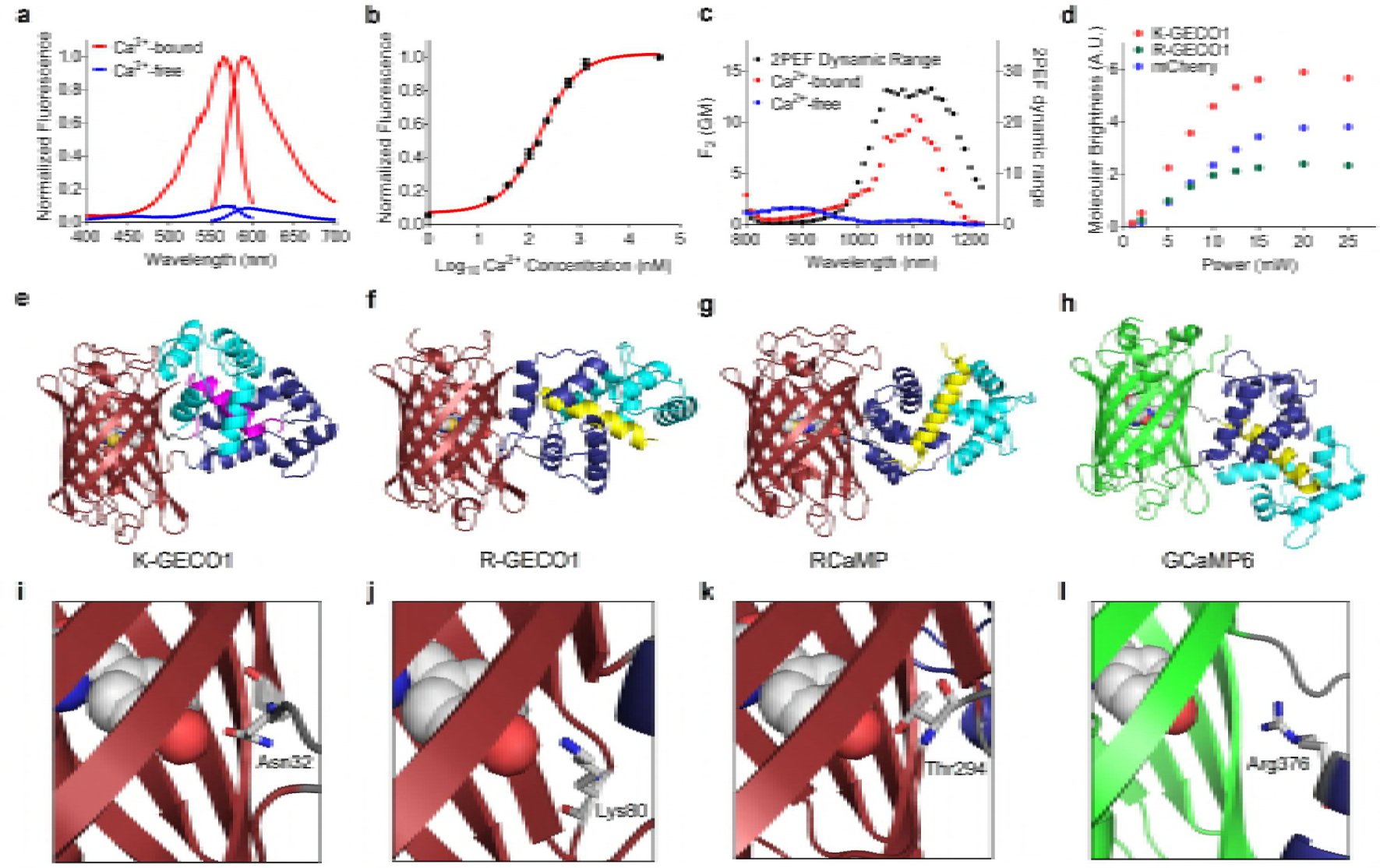
Characterization and structure of K-GECO1. **(a)** Fluorescence excitation and emission profile of K-GECO1 in the presence and absence of Ca^2+^. **(b)** Ca^2+^ titration curve of K-GECO1. **(c)** K-GECO1 effective two-photon fluorescence excitation spectra in Ca^2+^-saturated (red symbols) and Ca^2+^-free (blue symbols) states. Ratio of the K-GECO1 two-photon excitation fluorescence Ca^2+^-saturated/Ca^2+^-free signals as a function of wavelength (black symbols, plotted on right y-axis). **(d)** Two-photon molecular brightness of K-GECO1, R-GECO1 and mCherry with excitation at 1060 nm using various laser power. Overall protein structures of GECIs **(e-g)**, K-GECO1 (PDB: 5UKG) **(e)**, R-GECO1 (PDB: 4I2Y^19^) **(f)**, RCaMP (PDB: 3U0K^19^) **(g)**, and GCaMP6 (PDB: 3WLD^60^) **(h)**, with ckkap colored in magenta, RS20 in yellow, CaM N-lobe in dark blue, and CaM C-lobe in cyan. Zoom-in view of the interactions between key residues and the chromophore**(i-l)**, K-GECO1 **(i)**, R-GECO1 **(j)**, RCaMP **(k)**, and GCaMP6 **(l)**.

K-GECO1 displayed moderate photoactivation when illuminated with either a 405 nm or 488 nm laser, in both the Ca^2+^-free and Ca^2+^-bound states. For Ca^2+^-bound K-GECO1, illuminating with 405 nm (1.76 W/cm^2^) or 488 nm (6.13 W/cm^2^) laser light for 1 s resulted in a ~20% increase in fluorescence as detected using 561 nm illumination. For Ca^2+^-free K-GECO1, 1 s of 405 nm (1.76 W/cm^2^) or 488 nm (6.13 W/cm^2^) laser light also resulted in a ~20% increase in fluorescence **(Supplementary Fig. 2a)**. Consistent with previous reports^19,21^, we observed a more pronounced photoactivation with R-GECO1, but not RCaMP1h, under similar illumination conditions **(Supplementary Fig. 2b, c, d)**.

K-GECO1 shows a strong 2-photon excitation peak at approximately 1100 nm (**Fig. 2c**) in the Ca^2+^-bound state. A ~25-fold maximal increase of fluorescence signal, using 2-photon excitation in the excitation region from 1050 to 1150 nm, occurs upon binding Ca^2+^ (**Fig. 2c**). The peak 2-photon molecular brightness of K-GECO1 was compared with R-GECO1, using mCherry as a standard with 1060 nm excitation. The peak 2-photon molecular brightness, defined as the maximum detected fluorescence count rate per emitting molecule^35^, was obtained from the average fluorescence count rate and the average number of emitting molecules in the beam as determined by fluorescence correlation spectroscopy (FCS). Using this approach, K-GECO1 was found to be approximately 1.5-fold brighter than mCherry and over 2-fold brighter than R-GECO1 (**Fig. 2d**), which is consistent with the comparison of one-photon brightness for the Ca^2+^-bound state (**Supplementary Table 1**).

### Crystal structure of K-GECO1

To gain insight into the molecular mechanism of K-GECO1 Ca^2+^ sensitivity and to assist future protein engineering efforts, we determined the x-ray crystal structure of K-GECO1 in the Ca^2+^-bound form. The structure was determined to 2.36 Å resolution by molecular replacement (**Fig. 2e**, **Supplementary Table 2**). The crystal structure reveals the distinctive features of the ckkap/CaM complex in K-GECO1 (and presumably in other ckkap-based GECIs) relative to other RS20/CaM-based GECIs including R-GECO1 (**Fig. 2f**), RCaMP (**Fig. 2g**), and GCaMP6 (**Fig. 2h**). The major difference is that the binding orientation of the ckkap peptide to the CaM domain is opposite to that of RS20 to CaM^36,37^. Another difference is that the RS20 peptide consists entirely of an α-helix in the CaM-binding region, whereas the CaM-binding region of ckkap consists of both an α-helical segment as well as a hairpin-like loop structure at its C-terminus^34^.

Examination of the molecular interactions between the protein and the chromophore at the circular permutation site provides insights into the mechanism of Ca^2+^-dependent fluorescence modulation. The side chain of Asn32 of linker1 is in direct hydrogen bonding with the phenolate oxygen of the chromophore (**Fig. 2i**), and is positioned similarly to Ser143 of FusionRed, which engages in a similar interaction with the chromophore^9^. We reason that Asn32 plays a critical role in communicating the Ca^2+^-dependent conformational change in the ckkap/CaM domain to the chromophore in the cpRFP domain. Lys79 of R-GECO1 (**Fig. 2j**), Thr243 of RCaMP1h (**Fig. 2k**), and Arg376 of GCaMP6 (**Fig. 2l**), are likely to have similar roles in their respective mechanisms of fluorescence modulation. Saturation mutagenesis of Asn32 of K-GECO1 resulted in a library of variants that all had dimmer fluorescence and/or smaller Ca^2+^-induced fluorescence intensity fold change. This results indicates that Asn is the optimal residue in this position.

### Performance of K-GECO1 in cultured cells

To demonstrate the utility of K-GECO1 for imaging of Ca^2+^ dynamics, we expressed it in cultured human cells, dissociated rat neurons, organotypic rat brain slices, zebrafish sensory neurons, and mouse primary visual cortex. We first recorded the response of K-GECO1 to changes in the cytoplasmic Ca^2+^ concentration in HeLa cells using established protocols (**Fig. 3a**)^38^. HeLa cells expressing K-GECO1 had maximum fluorescence intensity changes of 5.2 ± 1.1-fold (n = 44) on treatment with histamine, which is similar with to the 4.9 ± 1.9-fold (n = 22) response previously reported for R-GECO1 expressing HeLa cells^18^.

**Figure 3.**
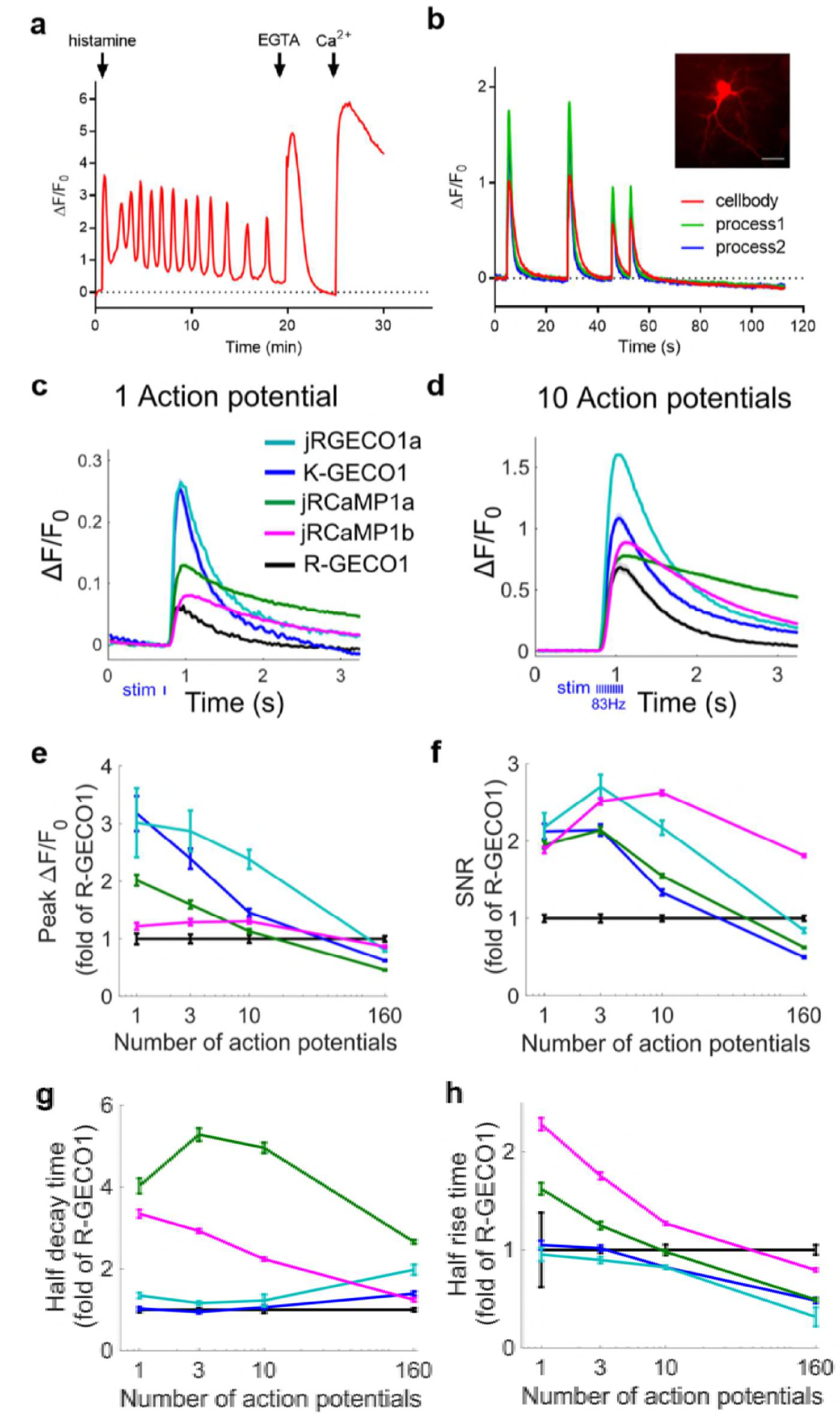
Performance of K-GECO1 in HeLa cells and cultured dissociated neurons. **(a)** Representative fluorescence time-course traces for HeLa cells expressing K-GECO1 with pharmacologically-induced Ca^2+^ changes. **(b)** Imaging of spontaneous Ca^2+^ oscillations in dissociated neurons expressing K-GECO1. Inset: fluorescence image of dissociated neurons expressing K-GECO1 (scale bar: 30 μm). **(c)** Average responses in response to one action potential (AP) for K-GECO1 compared with other red GECIs (same color code in panel c-h). **(d)** 10 APs response of red GECIs. Comparison of K-GECO1 and other red GECIs, as a function of number of APs **(e-h)**. Response amplitude, ΔF/F0 **(e)**. Signal-to-noise ratio (SNR) **(f)**. Half decay time (g). Half rise time **(h)**.

Next, we tested K-GECO1 in dissociated rat hippocampal neurons. The relatively low Ca^2+^ *K*_d_ of 165 nM for K-GECO1 is comparable to that of current best green GECI, GCaMP6s^17^, which has been highly optimized for detection of neuronal Ca^2+^ transients. Cultured dissociated neurons expressing K-GECO1 had fluorescence distributed throughout the cytosol and nucleus, and exhibited close to 2-fold maximum increases for spontaneous Ca^2+^ changes (**Fig. 3b**). We did not observe intracellular fluorescent punctate structures, as have been observed for R-GECO1 and its variants^22,27^, in the cell bodies of dissociated neurons expressing K-GECO1 (**Supplementary Fig. 3a,b**). We also did not observe noticeable photoactivation of K-GECO1 in neurons when illuminated with 0.5 W/cm^2^ 405 nm laser light. Under the same illumination conditions, R-GECO1 exhibited substantial photoactivation (**Supplementary Fig. 3c,d**). The absence of photoactivation for K-GECO1 under these conditions might be due to the relative low laser intensity (0.5 W/cm^2^) compared with the intensity (1.76 W/cm^2^) used for *in vitro* characterization.

To compare the performance of K-GECO1 with other red GECIs in dissociated neurons, we performed an automated imaging assay with field stimulation as previously described^17,24^. For a single action potential, K-GECO1 exhibited a similar response to jRGECO1a (**Fig. 3c**) and GCaMP6s^17^, two of the most sensitive indicators currently available. The peak ΔF/F**0** amplitude of K-GECO1 with 3 or more action potentials wassmaller than that of jRGECO1a, yet better than other red GECIs (**Fig. 3d, e**). In terms of signal-to-noise ratio (SNR), K-GECO1 had similar performance to jRGECO1a, but less than that of jRCaMPa/b (**Fig. 3f**). K-GECO1 exhibits fast kinetics, with has a half decay time that is faster than jRGECO1a and jRCaMP1a/b (**Fig. 3g**), and a half rise time that similar to jRGECO1a but faster than jRCaMP1a/b (**Fig. 3h**).

As our *in vitro* characterization indicated that K-GECO1 has less blue-light photoactivation than R-GECO1, we tested its performance in human induced pluripotent stem cell-derived cardiomyocytes (iPSC-CMs) in combination with Channelrhodopsin-2 (ChR2). As expected, transfected iPSC-CMs expressing K-GECO1 exhibited spontaneous Ca^2+^ oscillations (**Fig. 4a**). To compare photoactivation of K-GECO1 and R-GECO1 in iPSC-CMs, we illuminated transfected cells (GECI only, no ChR2) with 0.19 W/cm^2^ 470 nm LED light (**Fig. 4b, c**). Under these conditions, R-GECO1 exhibited a substantial photoactivation effect with a transient 200% increase in red fluorescence. Under the same illumination conditions, K-GECO1 had negligible change in red fluorescence. When we co-transfected iPSC-CMs with both K-GECO1 and ChR2, blue light stimulation reliably induced Ca^2+^ transients (**Fig. 4d**), demonstrating that the combination of K-GECO1 and ChR2 are suitable for all-optical excitation and imaging of iPSC-CMs.

**Figure 4.**
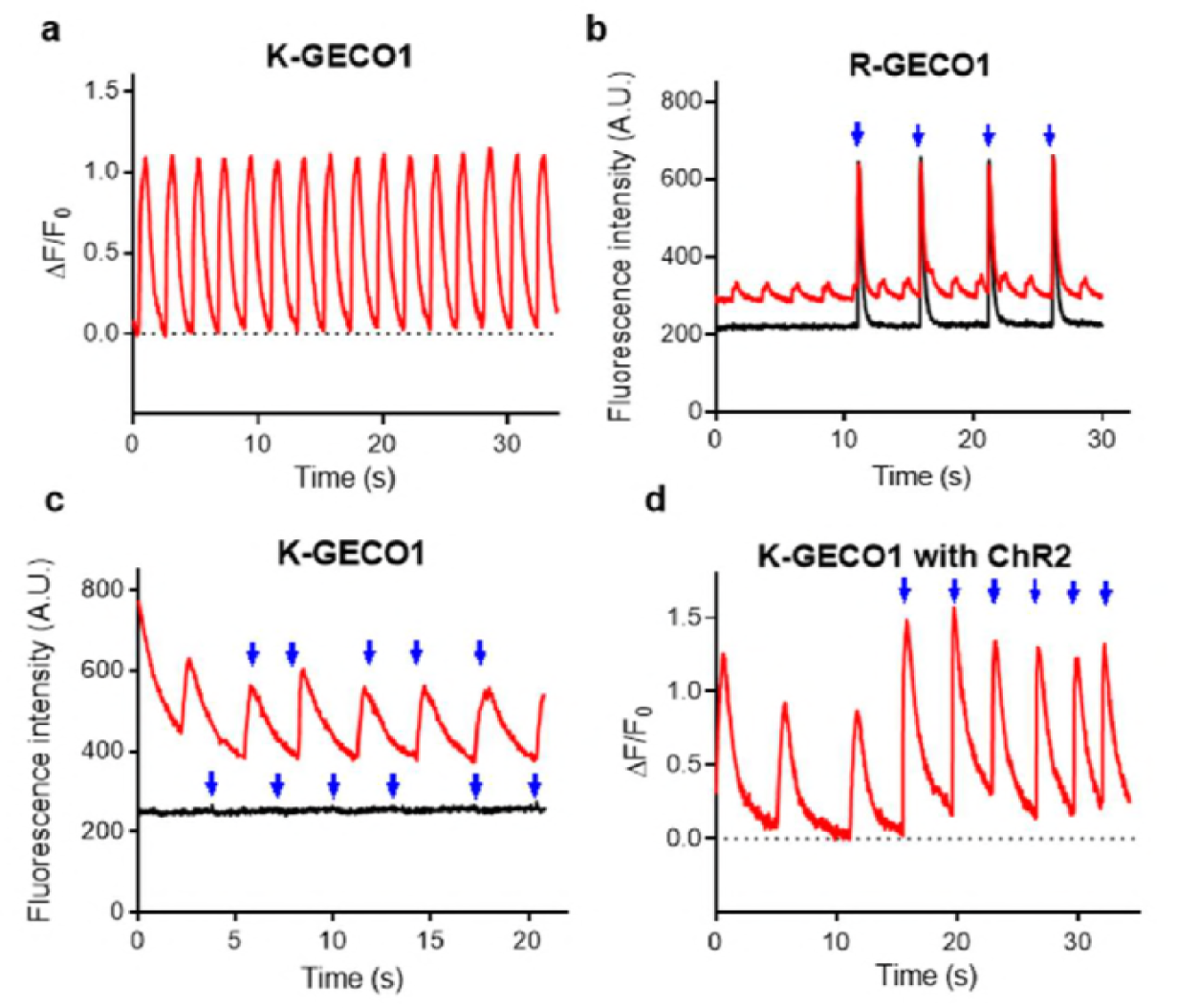
Performance of K-GECO1 in iPSC-CMs. (**a**) Representative time course of spontaneous Ca^2+^ oscillations in iPSC-CMs as imaged using K-GECO1. Photoactivation of R-GECO1 (**b**) and K-GECO1 (**c**) in iPSC-CMs. Cells with spontaneous activity are colored in red and cells with no spontaneous activity are colored in black. **(d)** Combined use of K-GECO1 with ChR2. Illumination with 150ms of 470 nm light is indicated by blue arrowheads.

### Performance of K-GECO1 in organotypic brain slices

We further tested the performance of K-GECO1 by expressing it in organotypic slices of the newborn rat ventromedial nucleus (VMN) of the hypothalamus. Expression of K-GECO1 enabled visualization of both neuronal cell bodies and processes (**Fig. 5a**). We investigated the performance of K-GECO1 under pharmacological stimulation by ATP (100 μM), which activates suramin-sensitive ATP-receptors and induces an influx of extracellular Ca^2+^, thus increasing cytosolic Ca^2+^ concentration^39^. Upon treatment with ATP, neurons expressing K-GECO1 underwent a mean increase in fluorescence intensity of 3.26 ± 0.18-fold (n = 21) (**Fig. 5b**).

**Figure 5.**
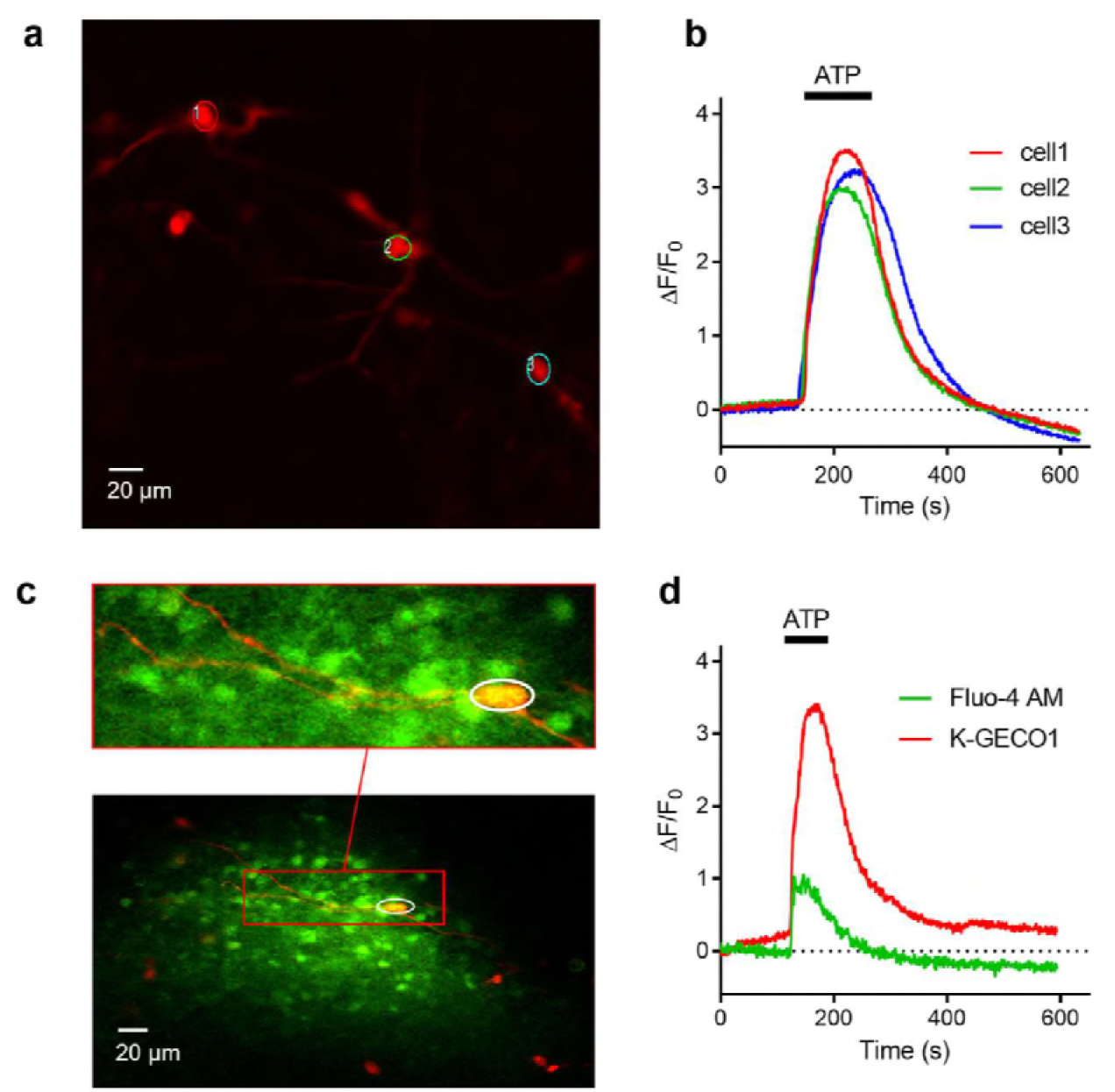
Performance of K-GECO1 in organotypic brain slice. **(a)** K-GECO1 labeling of the soma and dendrites of neurons in the ventromedial nucleus (VMN) of organotypically cultured newborn rat hypothalamus slices. (b) ATP-induced cytosolic Ca^2+^ rise in VMN neurons. (c) Fluo-4 AM loaded and K-GECO1 transfected into VMN slice. (d) ATP treatment caused a Ca^2+^ rise reported by both Fluo-4AM and K-GECO1. Schematic setup of the experiment. (**b**) Image of RB cells expressing K-GECO1 with ROI indicating cytoplasm. (**c**) K-GECO1 Ca^2+^ response to pulse stimuli in cytosol. (**d**) K-GECO1 Ca^2+^ response to pulse stimuli in nucleus. Fluorescence fold change of K-GECO1 (**e**) and jRGECO1a (**f**) under various numbers of pulses. Half decay time of K-GECO1 (**g**) and jRGECO1a (**h**) under various numbers of pulses.

To compare the performance of K-GECO1 with the small molecule-based green cytosolic Ca^2+^ indicator, Fluo-4AM, we loaded the dye into VMN neurons that were expressing K-GECO1 (**Fig. 5c**). When treated with ATP, these neurons (n=3) exhibited a 3.01 ±0.86-fold increase in K-GECO1 fluorescence, but only a 0.70 ± 0.12-fold increase in Fluo-4 fluorescence (**Fig. 5d**). In non-transfected cells stained with Fluo-4AM, we did not observe any crosstalk from Fluo-4AM into the red channel. Overall, K-GECO1 unravels robust responses to cytosolic Ca^2+^ concentration changes in neurons in organotypic brain slices.

### *In vivo* Ca^2+^ imaging with K-GECO1

To test K-GECO1 in zebrafish spinal cord sensory neurons *in vivo*, we transiently expressed K-GECO1 in Rohon Beard cells. Zebrafish Rohon-Beard (RB) cells have previously been used for *in vivo* GECI imaging and shown to fire a single spike in response to each electrical pulses to the skin^40^. Electrical stimulations were applied to trigger Ca^2+^ transients at 3 days post fertilization. 2-photon imaging with excitation at 1140 nm (**Fig. 6a**) revealed that K-GECO1 filled both the cytoplasm and nucleus *in vivo* in zebrafish RB neurons (**Fig. 6b**). Cytoplasmic K-GECO1 exhibited ~40% fluorescence intensity increase to Ca^2+^ transients triggered by a single pulse stimulus (**Fig. 6c**). When the RB neurons were stimulated with 5 to 20 repetitive stimuli, 50-100% increases in K-GECO1 fluorescence were observed (**Fig. 6d**). As expected, the fluorescence response in the nucleus was diminished with respect to the response in the cytosol, and exhibited a slower recovery to baseline (**Fig. 6c, d**). Compared to the optimized red fluorescent indicator jRGECO1a, K-GECO1 showed decreased sensitivity in zebrafish in terms of stimulus-evoked fluorescence change (**Fig. 6e, f**), whereas the half decay time was comparable (**Fig. 6g, h**). Consistent with the results from dissociated neurons, even distribution of the K-GECO1 red fluorescence in RB cells were observed in zebrafish neurons *in vivo* (**Supplementary Fig. 4a,b**), while jRGECO1 exhibited fluorescence accumulations (**Supplementary Fig. 4c**).

**Figure 6.**
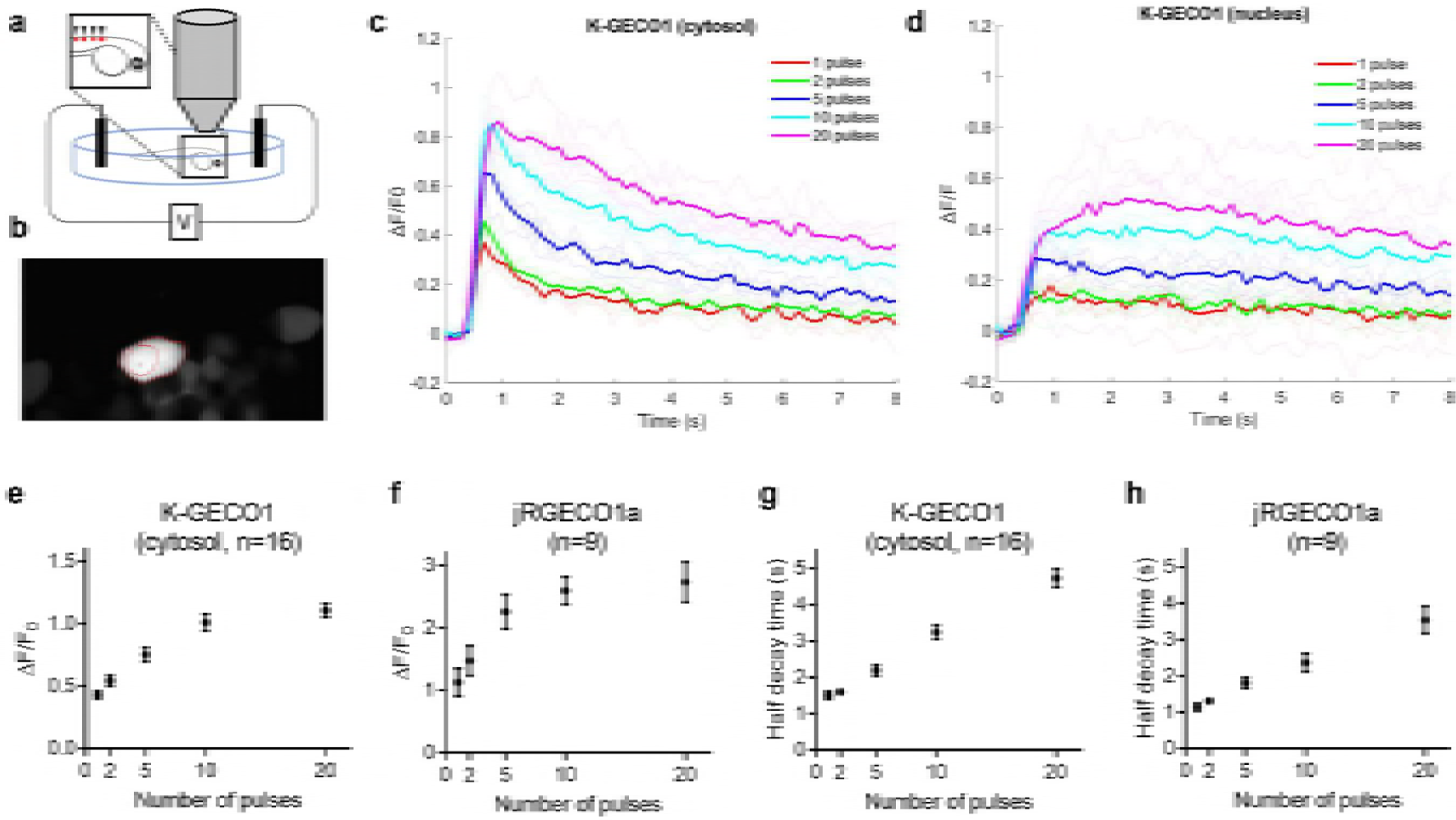
In vivo imaging of K-GECO in zebrafish Rohon-Beard (RB) cells. **(a)** Schematic setup of the experiment. **(b)** Image of RB cells expressing K-GECO1 with ROI indicating cytoplasm. **(c)** K-GECO1 Ca^2+^ response to pulse stimuli in cytosol. **(d)** K-GECO1 Ca^2+^ response to pulse stimuli in nucleus. Fluorescence fold change of KGECO1 **(e)** and jRGECO1a **(f)** under various numbers of pulses. Half decay time of K-GECO1 **(g)** and jRGECO1a **(h)** under various numbers of pulses.

To evaluate K-GECO1 in the mouse primary visual cortex (V1) *in vivo,* V1 neurons were infected with adeno-associated virus (AAV) expressing nuclear export signal (NES) tagged K-GECO1 under the human synapsin1 promoter (AAV-SYN1-NES-K-GECO1). The majority of V1 neurons can be driven to fire action potentials in response to drifting gratings. Eight-direction moving grating visual stimuli were presented to the contralateral eye (**Fig. 7a**). K-GECO1 expressing L2/3 neurons exhibited cytoplasmic red fluorescence (**Fig. 7b**), and 2-photon imaging revealed visual stimulus-evoked fluorescence transients in subsets of neurons (**Fig. 7c**). We compared the performance of K-GECO1 with other red GECIs using previously established metrics^17,24^. In terms of the fraction of neurons detected as responsive in the visual cortex, K-GECO1 is higher than RCaMP1h, but lower than R-GECO1 and other optimized red indicators (**Fig. 7d**). The mean ΔF/F**0** at the preferred visual stimulus is reflective of indicator sensitivity. By this metric, K-GECO1 has sensitivity that is comparable to R-GECO1 and jRCaMP1a, but less than jRGECO1a (**Fig. 7e**). Lysosomal accumulation was previously observed in mouse V1 neurons labeled with jRGECO1a, but not in the ones with jRCaMP1a/b^24^. Fixed brain tissue sections, prepared as previously reported for jRGECO1a and jRCaMP1a/b^24^, revealed no signs of lysosomal accumulation in K-GECO1 expressing V1 neurons (**Supplementary Fig. 5a**). As with both jRGECO1a and jRCaMP1a/b, *in vivo* functional imaging of K-GECO1 did exhibit fluorescent clump-like structures (**Supplementary Fig. 5b**), yet these structures were not observed in fixed sections of the same tissue. We are currently unable to explain this discrepancy. Overall, the results demonstrate that K-GECO1 can be used to report physiological Ca^2+^ changes in neurons *in vivo* with performance that matches or surpasses that of other first-generation red fluorescent Ca^2+^ indicators.

**Figure 7.**
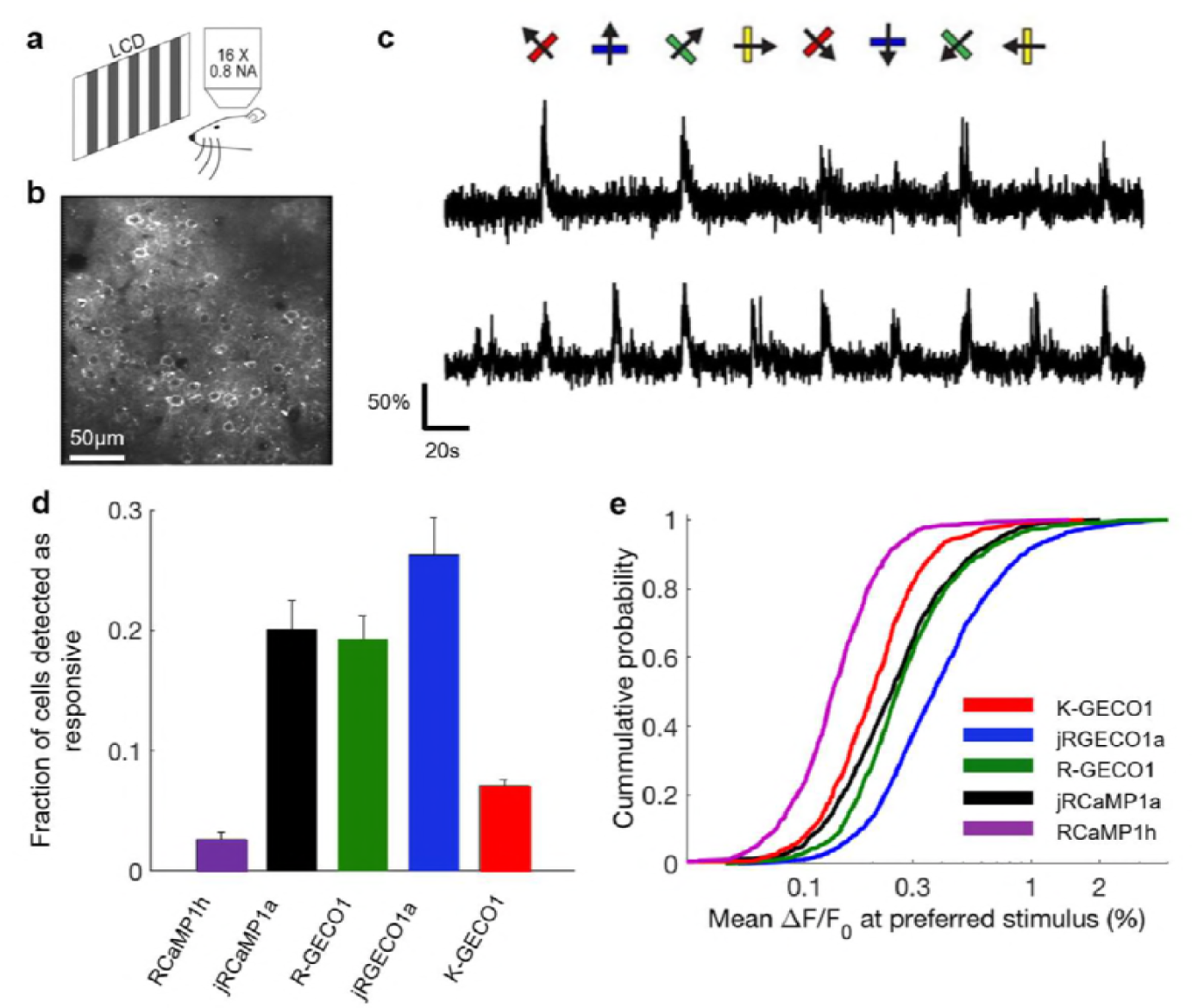
*In vivo* imaging of K-GECO1 in mouse V1 neuron. **(A)** Schematic setup of the experiment. (**b**) Image of V1 L2/3 cells expressing K-GECO1. (**c**) Example traces from neurons expressing K-GECO1. Directions of grating motion are indicated above the traces. (**d**) Fraction of cells detected as responding to visual stimulus of K-GECO1 compared with previously reported values^24^ from other red GECIs. (**e**) Distribution of ΔF/F**0** amplitude for the preferred stimulus of K-GECO1 compared with previously reported values^24^ from other red GECIs.

## Discussion

Although green fluorescent GECIs are currently the most highly effective tools for *in vivo* visualization of neuronal signaling, we anticipate that they will one day be made redundant by red fluorescent GECIs due to the inherent advantages associated with longer wavelength fluorescence. The transmittance of tissue increases as wavelength increases, so red fluorescent GECIs will enable imaging of neuronal activity deeper into brain tissue than is possible with green fluorescent GECIs, assuming all other properties are equivalent^24,41^. In addition, red fluorescent GECIs enable multiparameter imaging in conjunction with green fluorescent indicators, and facilitate simultaneous imaging and optical activation when used in conjunction blue light activatable optogenetic actuators such as channelrhodopsin-2 (Ref. 42). However, as widely recognized^13,19,22,24^, red GECIs currently suffer from a number of limitations compared to the most highly optimized green GECIs (i.e., GCaMP6)^17^. These limitations include decreased sensitivity for RCaMP variants and complicated photophysics and lysosomal accumulation for R-GECO variants. As both green and red GECIs have analogous designs and contain identical Ca^2+^-binding domains, these undesirable characteristics are related to the RFP scaffold used to generate red GECIs.

In an effort to overcome the limitations associated with current RFP scaffolds, we turned our attention to the eqFP578-derived lineage of monomeric RFPs (i.e., mKate and its derivatives)^7–9^, which tend to give bright and evenly-distributed fluorescence when expressed in the neurons in transgenic mice^30^. Using semi-rational design and directed evolution we developed a new red fluorescent Ca^2+^ indicator, K-GECO1, based on a mKate variant FusionRed^9^. We anticipated that K-GECO1 would retain the favorable traits associated with its starting template RFP. We have found this expectation to be generally true, as we did not observe lysosomal aggregation in dissociated rat neurons, or zebrafish neurons, or fixed mouse brain tissue expressing K-GECO1. Some fluorescent punctate-like structures were observed during *in vivo* functional imaging.

The other distinctive feature of K-GECO1 is the use of the ckkap peptide as the CaM binding partner for the Ca^2+^-binding motif. Consistent with previous reports^23,34^, the ckkap/CaM motif yielded a lower apparent *K*_d_ for Ca^2+^ and faster kinetics (relative to RS20/CaM), and an apparent Hill coefficient close to 1. These characteristics should enable more sensitive detection of Ca^2+^ dynamic at physiological ranges, as is evident from K-GECO1’s large single action potential fluorescence response amplitude. With a Hill coefficient close to 1, K-GECO1 should provide a more linear Ca^2+\^ response following multiple stimuli.

The x-ray crystal structure of K-GECO1 suggests that the indicator has a “self-contained” fluorescence modulation mechanism, similar to that proposed for R-GECO1 (Ref. 22,29). Unlike GCaMP, in which the fluorescence modulation mechanism is dependent on the interactions with a residue of CaM^43^ (**Fig. 2l**), the K-GECO1 Ca^2+^-bound state is likely stabilized by the hydrogen bonding between the phenolate group of chromophore and linker1 residue Asn32 (**Fig. 2i**). This makes the cpFusionRed protein in K-GECO1 a potentially useful template as a signal transduction domain to be combined with other binding domains for development of new types of red fluorescent indicators. The crystal structure also reveals that the ckkap/CaM motif in K-GECO1 has a reversed binding orientation for CaM when compared with the RS20/CaM binding patterns in R-GECO1, RCaMP, and GCaMP6 (**Fig. 2e-h**). These results indicate that the GCaMP-like design is versatile enough to tolerate different peptide conformations and CaM orientations, and that exploring a wider range of CaM binding partners is likely to lead to GECIs with new and improved properties.

First generation red GECIs, including mApple-based R-GECO1 and mRuby-based RCaMP1h, have been optimized using a neuron screening platform^24,44^, resulting in jRGECO1a and jRCaMP1a/b with greatly improved *in vivo* performance for detection of action potentials. Although K-GECO1 is a first generation red GECI, it already provides performance that, by some criteria, is comparable to second generation red GECIs. Specifically, K-GECO1 has a fluorescent response to single action potentials that is similar to that of jRGECO1a (and superior to jRCaMP1a/b) and faster dissociation kinetics than either jRGECO1a or jRCaMP1a/b. However, by other criteria, K-GECO1 will require further optimization in order to match the performance of second generation red GECIs. For example, K-GECO1 does not provide the same level of *in vivo* sensitivity as the highly optimized jRGECO1a. In addition, K-GECO1 showed some blue light-dependent photoactivation during *in vitro* characterization, though less than R-GECO1. The photoactivation of K-GECO1 was not detectable in our characterization in cultured dissociated neurons (**Supplementary Fig. 3c**) or in iPSC-CMs (**Fig.4c**), suggesting that it is more suitable than R-GECO1 for use with blue/cyan excitable optogenetic actuators.

In summary, we have demonstrated the utility of K-GECO1 in various cell types including HeLa cells, dissociated neurons, iPSC-CMs, neurons in organotypic rat brain slices, zebrafish RB cells, and mouse V1 neurons *in vivo*. Though not yet ideal by all criteria, K-GECO1 represents a step forward in the development of red GECIs. As with R-GECO1 and RCaMP1h, further optimization using a neuron-based screening approach is likely to yield K-GECO variants with much improved sensitivity and performance *in vivo*.

## Methods

### Protein engineering

The design of K-GECO is based on well-established GECI designs reported previously^18,32,45-47^. Initial construction of mKate2 and FusionRed-based Ca^2+^ indicators was done by overlapping assembly of the four DNA parts encoding the following protein fragments: the N-terminal (1-145) and C-terminal (146-223) parts of mKate2 or FusionRed, the RS20 peptide, and the CaM of R-GECO1. The fragments were PCR amplified from mKate2, FusionRed (a kind gift from Michael Davidson), and R-GECO1 DNA. Overlap region and restriction sites were encoded in the primers. DNA encoding ckkap was synthesized by IDT. Purified PCR products were pooled and assembled in an overlapping PCR reaction. The resulting assembled PCR product was purified, digested with *XhoI* and *HindIII* (Thermo Fisher Scientific), and then ligated into a similarly digested pBAD/His B vector (Thermo Fisher Scientific). The ligation product was transformed into electrocompetent *E. coli* strain DH10B cells. Plasmids were purified with the GeneJET miniprep kit (Thermo Fisher Scientific) and then sequenced using the BigDye Terminator Cycle Sequencing kit (Thermo Fisher Scientific).

For construction of random mutagenesis libraries, Error-prone PCR (EP-PCR) amplifications were performed. The EP-PCR products were digested with *XhoI* and *HindiIII*, and then ligated into a similarly digested pBAD/His B vector (Thermo Fisher Scientific). For site-directed mutagenesis and saturation mutagenesis libraries construction, QuikChange site-directed mutagenesis Lightning Single or Multi kit (Agilent Technologies) was used according to the manufacturer's instructions. The resulted variant libraries were transformed into electrocompetent *E. coli* strain DH10B cells and incubated overnight at 37°C on 10 cm Petri dishes with LB-agar supplemented with 400 g/mL ampicillin (Sigma) and 0.02% (wt/vol) L-arabinose (Alfa Aesar).

A custom imaging system was used for screening K-GECOs on plate with *E. coli* colonies expressing the variants^48^. When screening, fluorescence images of E. *coli* colonies was taken for each Petri dish with an excitation filter of 542/27 nm and an emission filter of 609/57 nm. The colonies with the highest fluorescence intensity in each image were then picked and cultured in 4 mL liquid LB medium with 100 μg/ml ampicillin and 0.02% L-arabinose at 37°C overnight. Proteins were then extracted using B-PER reagents (Thermo Fisher Scientific) from the liquid culture. The protein extraction was used for a secondary screen of Ca^2+^-induced response test using Ca^2+^-free buffer (30 mM MOPS, 100 mM KCl, and 10 mM EGTA at pH 7.2) and Ca^2+^-buffer (30 mM MOPS, 100 mM KCl, and 10 mM Ca-EGTA at pH 7.2) in a Safire2 fluorescence microplate reader (Tecan).

### *In vitro* characterization

To purify K-GECO variants for *in vitro* characterization, pBAD/His B plasmid encoding the variant of interest was used to transform electrocompetent *E. coli* DH10B cells and then plated on LB agar plate with ampicillin (400 μg/mL). Single colonies were picked and inoculated into 5 mL LB medium supplemented with 100 g/mL ampicillin. Bacterial subcultures were incubated overnight at 37°C. 5 mL of bacterial subculture was added into 500 mL of LB medium with 100 μg/mL ampicillin. The cultures were incubated at 37°C to an OD of 0.6. Following induction with L-arabinose to a final concentration of 0.02% (wt/vol), the cultures were then incubated at 20°C overnight. Bacteria were harvested by centrifugation at 4,000 g for 10 min, resuspended in 30 mM Tris-HCl buffer (pH 7.4), lysed using French press, and then clarified by centrifugation at 13,000 g for 30 mins. Proteins were purified from the cell-free extract by Ni-NTA affinity chromatography (MCLAB). The buffer of purified proteins was exchanged into 10 mM MOPS, 100 mM KCl, pH 7.2. Absorption spectra were recorded on a DU-800 UV-visible spectrophotometer (Beckman) and fluorescence spectra were recorded on a Safire2 fluorescence plate reader (Tecan).

For quantum yield (QY) determination, the fluorescent protein mCherry was used as a standard. The detailed protocol has been described previously^18^. Briefly, the fluorescence emission spectra of each dilution of mCherry and K-GECO variants protein solution were recorded. The total fluorescence intensities were obtained by integration. Integrated fluorescence intensity versus absorbance was plotted for both mCherry and K-GECOs. QY was determined from the slopes of mCherry and K-GECOs. Extinction coefficient (EC) was determined by first measuring the absorption spectrum of K-GECO variants in Ca^2+^-free buffer and Ca^2+^-buffer. Following alkaline denaturation, the absorption was measured. With the assumption that the denatured chromophore has an EC of 44,000 M^−1^cm^−1^ at 446 nm, the protein concentration was determined. EC of K-GECO variants were calculated by dividing the peak absorbance maximum by the concentration of protein.

For the Ca^2+^ *K*_d_ determination, purified protein solution was diluted into a series of buffers which were prepared by mixing Ca^2+^-buffer and Ca^2+^-free buffer with free Ca^2+^ concentration ranges from 0 nM to 3,900 nM. The fluorescence intensity of K-GECO variants in each solution was measured and subsequently plotted as a function of Ca^2+^ concentration. Data was fit to the Hill equation to obtain *K*_d_ and the apparent Hill coefficient.

Two-photon excitation spectra and cross sections were measured as previously reported^49^, with the following adjustments. For the two-photon excited (2PE) spectra, fluorescence was collected through a 694/SP filter for K-GECO1 (Semrock). To correct for wavelength-to-wavelength variations in the laser parameters, a correction function using Rhodamine B in MeOH and its known 2PE spectrum was applied^50^. Two-photon cross sections were measured at 1100 nm for K-GECO1, with Rhodamine B in MeOH as a reference standard. The fluorescence for cross sections were collected through a narrow bandpass filter, 589/15 (Semrock), and differential quantum efficiencies were obtained at 582 nm with a PC1 ISS spectrofluorimeter (this wavelength corresponded to the bandpass center of the above filter when used in the MOM Sutter Instruments microscope due to its tilted position). Since the filter (694/SP) used for the 2PE spectra measurements covers the fluorescence of both the neutral and anionic forms of the chromophore, the spectrum of a particular Ca^2+^ state of a protein represents a combination of the unique 2PE spectra of the neutral and anionic forms, weighted to their relative concentrations (ρ, concentration of one form divided by the total chromophore concentration) and quantum yields. The y-axis of the total 2PE spectrum is defined by F_2_ (λ)= σ_2,N_(λ) φN ρN + σ_2,A_(λ) φA ρA, where σ_2_(λ) is the wavelength-dependent 2-photon cross section and φ is the fluorescence quantum yield of the corresponding form (N for neutral or A for anionic in the subscript). At the wavelengths used to measure the cross sections (1060 and 1100 nm), σ_2,N_ is assumed to be zero, and φA and ρA were independently measured to give a value for F_2_ (GM). The relative concentrations of the neutral and anionic forms were found by measuring the absolute extinction coefficients of each respective form in the Ca^2+^-free and the Ca^2+^-bound states. These differ from the effective extinction coefficients reported in Supplementary Table 1, which are weighted by the relative concentrations of both forms of the chromophore.

For FCS measurement of 2-photon molecular brightness, dilute protein (50-200 nM) solutions in Ca^2+^ buffer (30 mM MOPS, 100 mM KCl, 10 mM CaEGTA, pH 7.2) were excited at 1060 nm at laser powers from 1 - 25 mW for 200 second. At each laser power, fluorescence was recorded by the avalanche photodiode (APD) and fed to an Flex03LQ autocorrelator (Correlator.com). The measured autocorrelation curve was fit to a simple diffusion model with a custom Matlab program^35^ to determine the average number of excited molecules *<N>* in the excitation volume. The 2-photon molecular brightness (ε) at each laser power was calculated as the average rate of fluorescence <*F*> per emitting molecule <*N*>, defined as *ε* = *<F>/<N>* in kilocounts per second per molecule (kcpsm). As a function of laser power, the molecular brightness initially increases as the square of the laser power, then levels off and decreases due to photobleaching or saturation of the protein chromophore in the excitation volume. The maximum or peak brightness achieved, *<emax>,* represents a proxy for the photostability of a fluorophore.

For measuring photoswitching of K-GECO1, R-GECO1, and RCaMP1h *in vitro,* the purified protein in Ca^2+^ buffer (30 mM MOPS, 100 mM KCl, 10 mM CaEGTA, pH 7.2) or EGTA buffer (30 mM MOPS, 100 mM KCl, 10 mM EGTA, pH 7.2) were made into aqueous droplets with octanol in 1:9 ratio and mounted on a presilanized coverslip. A single droplet was focused under the AxioImager microscope (Zeiss) with a 20x 0.8 NA objective and photoswitched by different laser excitation of 561 nm, 405 nm, and 488 nm. Fluorescence emission was detected using a SPCM-AQRH14 fiber coupled avalanche photodiode (Pacer).

### Protein crystallography

K-GECO1 DNA was cloned into pRSET-A with short N-terminal hexahistidine purification tag <u>(MHHHHHHGSV</u>KLIP..., tag underlined). K-GECO1 was expressed in T7 Express *E. coli* cells (New England Biolabs) for 36 h in autoinduction medium^51^ supplemented with 100 mg/L ampicillin. *E. coli* pellets were lysed in B-PER (Thermo Fisher Scientific) supplemented with 1 mg/mL lysozyme followed by sonication. Insoluble cell debris was removed from the lysate by centrifugation for 20 min at 25,000 ×g, and soluble K-GECO1 protein was purified by immobilized metal affinity chromatography with nickel-charged Profinity resin (Bio-Rad), washing with 10 mM imidazole and eluting with 100 mM imidazole in Tris-buffered saline. K-GECO1 was further purified by size exclusion chromatography using a Superdex 200 column (GE Healthcare Life Sciences) with 10 mM Tris, 100 mM NaCl, pH 8.0 as the mobile phase. Purified K-GECO was concentrated to 10 mg/mL for crystallization using centrifugal concentrators (Sartorius Vivaspin, 10,000 MWCO). Purified K-GECO1 protein at 10 mg/mL in 10 mM Tris, 100 mM NaCl, pH 8.0 was mixed with an equal volume of a precipitant solution containing 100 mM BIS-TRIS, 20% w/v polyethylene glycol monomethyl ether 5,000, pH 6.5 at room temperature in a sitting-drop vapor diffusion crystallization tray (Hampton Research). Crystals were cryoprotected in the precipitant solution supplemented with 25% ethylene glycol. X-ray diffraction data were collected at 100 K on beamline 8.2.1 of the Advanced Light Source. Diffraction data were processed using the HKL software package^52^. The structure was solved by molecular replacement using Phaser^53^, searching first for two copies of the fluorescent protein domain fragment using a single molecule of mKate (PDB ID 3BXB) as the search model, followed by two copies each of the separated N- and C-terminal lobes of the Ca^2+^-bound calmodulin domain using fragments of PDB ID 3SG3. Iterative model building in Coot^54^ and refinement in Refmac^55^ produced the K-GECO1 model, with two copies of K-GECO1 in the asymmetric unit. The K-GECO1 model was deposited at the PDB with the accession code of 5UKG.

### Cell culture and imaging

For K-GECO variants characterization in HeLa cells, the cells were maintained in Dulbecco’s modified Eagle medium (DMEM) supplemented with 10% fetal bovine serum (FBS, Thermo Fisher Scientific), Penicillin-Streptomycin (Thermo Fisher Scientific), GlutaMAX (Thermo Fisher Scientific) at 37°C with 5% CO_2_. To construct the mammalian expression plasmid, pcDNA3.1(+) and the K-GECO variant were both digested with *XhoI* and *HindIII,* and the digested plasmid backbone and insert were purified by gel electrophoresis, followed by ligation and sequencing confirmation. Transient transfections of pcDNA3.1(+)-K-GECO plasmids were performed using Lipofectamine 2000 (Thermo Fisher Scientific). HeLa cells (60-70% confluency) on 35 mm glass bottom dishes (In vitro scientific) were transfected with 1 μg of plasmid DNA, using Lipofectamine 2000 (Thermo Fisher Scientific) according to the manufacturer's instructions. The cells were imaged 24 h after the transfection. Immediately prior to imaging, cells were washed twice with Hanks balanced salt solution (HBSS) and then 1 mL of 20 mM HEPES buffered HBSS (HHBSS) was added. Cell imaging was performed with an inverted Eclipse Ti (Nikon). The AquaCosmos software package (Hamamatsu) was used for automated microscope and camera control. Cells were imaged with a 20× objective lens. For imaging of histamine-induced Ca^2+^ dynamics, cells were imaged with a 200 ms exposure acquired every 5 s for a duration of 30 min. Approximately 60 s after the start of the experiment, histamine (10 μL) was added to a final concentration of 5 mM. The oscillation was imaged for 20 min, EGTA/ionomycin (40 μL) in HHBSS was added to a final concentration of 2 mM EGTA and 5 μM of ionomycin. After 5 min, Ca^2+^/ionomycin (40 μL) in Ca^2+^ and Mg^2+^ free HHBSS was added to a final concentration of 5 mM Ca^2+^ and 5 μM of ionomycin.

For K-GECO variants characterization in cultured dissociated neurons, the procedure was done as previously reported^29^. Dissociated E18 Sprague Dawley hippocampal cells were purchased from BrainBits LLC. The cells were grown on 35 mm glass bottom dish (In Vitro Scientific) containing NbActiv4 medium (BrainBits LLC) supplemented with 2% FBS, penicillin-G potassium salt (50 units/ml), and streptomycin sulfate (50 mg/ml). Half of the culture media was replaced every 4-5 days. Cells were transfected on day 8 using Lipofectamine 2000 (Thermo Fisher Scientific) following the manufacturer's instructions with the following modifications. Briefly, 1-2 μg of plasmid DNA and 4 μl of Lipofectamine 2000 (Thermo Fisher Scientific) were added to 100 μl of NbActive4 medium to make the transfection medium and incubated at room temperature for 10-15 min. Half of the culture medium (1 ml) from each neuron dish was taken out and combined with an equal volume of fresh NbActiv4 medium (supplemented with 2% FBS, penicillin-G potassium salt, and streptomycin sulfate) to make a 1:1 mixture and incubated at 37°C and 5% CO_2_. 1 ml of fresh conditioned (at 37°C and 5% CO_2_) NbActiv4 medium was added to each neuron dish. After the addition of transfection medium, the neuron dishes were incubated for 2-3 h at 37°C in a CO_2_ incubator. The medium was then replaced using the conditioned 1:1 mixture medium prepared previously. The cells were then incubated for 48-72 h at 37°C in a CO_2_ incubator before imaging. Fluorescence imaging was performed in HHBSS. Fluorescence imaging was performed on an inverted Nikon Eclipse Ti-E microscope equipped with a 200 W metal halide lamp (PRIOR Lumen), 60x oil objectives (numerical aperture, NA = 1.4; Nikon), a 16-bit QuantEM 512SC electron-multiplying CCD camera (Photometrics), and a TRITC/Cy3 filter set (545/30 nm excitation, 620/60 nm emission, and a 570LP dichroic mirror, Chroma). For time-lapse imaging, neurons were imaged at 100 Hz imaging frequency with 4×4 binning. For photoactivation comparison, cells expressing K-GECO1 and R-GECO1 were stimulated with pulses of blue laser light (405 nm, 5 mW/mm^2^).

For comparison of K-GECO1 and Red GECIs in stimulated cultured neuron cells, the procedure was done as previously reported^24^. Briefly, red GECIs were expressed after electroporation into rat primary hippocampal neurons (P0) using Nucleofector system (Lonza). For stimulation, action potentials were evoked by field stimulation. TxRed (540-580 nm excitation, 593-668 nm emission, and 585 nm long pass dichroic mirror) filter set was used for illumination. Responses were quantified for each cell as change in fluorescence divided by baseline fluorescence before stimulation. Signal-to-noise ratio (SNR) was quantified as the peak fluorescence signal over baseline, divided by the standard deviation of fluorescence signal before the stimulation.

iPSC-CMs were purchased from Axol Bioscience. Cells were plated in two wells of a 6-well plate and cultured for 4 days in Cardiomyocyte Maintenance Medium (Axol Bioscience) to 60-80% confluency. Cells then were then transferred to Fibronectin-coated (1%) coverslips and imaged in Tyrode’s buffer. Cells were transfected using transfection reagent Lipofectamine 2000 (Invitrogen). An inverted microscope (Zeiss) equipped with a NA 1.4, 63× objective lens (Zeiss) and a pE-4000 multi-wavelength LED light source (CoolLED) was used. Blue (470 nm) and green (550 nm) excitation were used to illuminate ChR2-EYFP and red GECIs, respectively. The GFP filter set (excitation 480/10 nm, 495 nm long pass dichroic mirror, emission 525/50 nm) and the RFP filter set (excitation 545/30, 565 nm long pass dichroic mirror, emission 620/60 nm) was used to visualize ChR2-EYFP and K-GECO or R-GECO, respectively. Optical stimulation was achieved with the 470 nm LED light at a power density of 0.19 W/cm^2^ and a pulse duration of 150 ms. Fluorescence signals were recorded using an ORCA-Flash4.0LT sCMOS camera (Hamamatsu) controlled by ImageJ^56^.

### Organotypic hypothalamic rat brain slice imaging

For the preparation of organotypic brain slices, experiments were done on neonatal rat coronal brain slices containing the ventromedial nucleus (VMN) of the hypothalamus. All procedures were carried out in compliance with the guidelines of the Canadian Council for Animal Care and with the approval of the University of Alberta Health Animal Care and Use Committee for Health Sciences. In brief, postnatal day 0-1 old Sprague-Dawley rats were anesthetized with 2-3% isoflurane until the paw reflex disappeared. Following decerebration, the brain was isolated in ice-cold divalent cation-free HBSS (Thermo Fisher Scientific) with 1 mM CaCl_2_ and 1.3 mM MgSO_4_ added. The brain was glued caudal side down to a metal plate and serial sections of 400 μm thickness were made using a vibratome (Leica Microsystems). Sectioning was stopped when the third ventricle became visible and two VMN-containing slices of 250 μm thickness were cut. Individual slices were placed on a sterile 0.4-μm-pore-membrane cell culture insert (Millipore). The insert and slice were then transferred to a 35 mm diameter culture dish (Corning) containing 1.5 ml of NbActiv4 medium (BrainBits) supplemented with 5% FBS, penicillin-G potassium salt (50 units/ml), and streptomycin sulfate (50 μg/ml). Slices were cultured at 37°C in an incubator (Thermo Fisher Scientific) under gassing with 5% CO_2_.

For transfection of organotypic slices, after 8-10 days of organotypic slice culturing, the VMN areas were transfected with an electroporation technique previously described^47^. Specifically, the insert with the slice was placed on a platinum plate petri dish electrode (Bex Co Ltd) and electroporation buffer (HBSS with 1.5 mM MgCl_2_ and 10 mM D-glucose) was filled between the electrode and the membrane. Plasmids of pcDNA3.1-K-GECO1 were dissolved in electroporation buffer at a concentration of 1 μg/ml and 10 μl of this solution was added to just cover the slice. Then, a square platinum electrode (Bex Co Ltd) was placed directly above the slice. Five 25 V pulses (5 ms duration, interval 1 s) were applied twice (the second time with reversed polarity) using a pulse stimulator (Sequim) and an amplifier (Agilent). The electroporation buffer was replaced with supplemented NbActiv4 medium and slices were returned to the incubator.

For imaging of cytosolic Ca^2+^ dynamics using K-GECO1, an upright FV1000 confocal microscope equipped with FluoView software and a 20× XLUMPlanF1 water immersion objective (NA 1.0) was used (Olympus). The millipore insert containing the transfected brain slice was placed in a custom-made chamber and mechanically fixed with a platinum harp. The slices were then perfused at 31 °C with artificial cerebrospinal fluid (ACSF) containing (in mM) 120 NaCl, 3 KCl, 1 CaCl_2_, 1.3 MgSO_4_, 26 NaHCO_3_, 1.25 NaH_2_PO_4_ and 10 D-glucose (pH was adjusted to 7.4 by gassing with 95% O_2_ plus 5% CO_2_), at a flow rate of 5 ml/min using a peristaltic pump (Watson-Marlow). For single-colour confocal C_ai_ imaging, K-GECO-transfected VMN neurons were exposed to excitation with 543 nm laser light and emission was collected from 560 nm to 660 nm using a variable barrier filter. Images were acquired at ×1-3 digital zoom at a frame resolution (512 × 512) and with a 2 μs/pixel scanning rate resulting in image acquisition at 1.12 frames/s. For monitoring drug-evoked cytosolic Ca^2+^ rises approximately 60 s after the start of image acquisition, 100 μΜ adenosine triphosphate (ATP, Sigma-Aldrich) was added to the ACSF for 90 s. For comparing K-GECO1 signal with that of a chemical Ca^2+^ fluorescent dye, transfected slices were stained with the membrane-permeant (AM) variant of green Fluo-4 by focal application. In brief, 0.5 mM Fluo-4-AM was filled into a broken patch pipette with an outer diameter of ~10 μm and subsequently pressure-injected (25-50 mmHg) for 10 min^57,58^ at 30-50 μm depth into the slice in the vicinity of K-GECO1-transfected VMN neurons. This led to uniform staining of cells in a radius of 150-200 μm from the injection site. For dual-colour imaging of K-GECO1- and Fluo-4-based Ca^2+^ responses, double-labelled neurons were excited with a 488 nm laser and emission was simultaneously collected in two channels from 500 to 520 nm for Fluo-4 and 570 to 670 nm for K-GECO1 using variable barrier filters.

### Zebrafish spinal sensory neurons imaging

Mitfa^w2/w2^ roy^a9/a9^ (Casper) zebrafish were maintained under standard conditions at 28 □ and a 14:10 hr light:dark cycle. Embryos (1-2 cell stage) of Tg (elavl3:GAL4-VP16)^59^ were injected with 25 ng/μΙ DNA plasmids encoding the K-GECO variants under the control of the 10xUAS promoter, and 25 ng/μL Tol2 transposase mRNA diluted in E3 medium. Three day post-fertilization embryos showing expression in spinal sensory neurons (Rohon-Beard cells) were paralyzed by 5-min bath application of 1 mg/ml a-bungarotoxin (Sigma, 203980). Larvae were mounted on their side in a field stimulation chamber (Warner, RC-27NE2) with 1.5% low melting point agarose and imaged using a custom-built two-photon microscope equipped with a resonant scanner. The light source was an Insight DS Dual femtosecond laser (Spectra-Physics) running at 1140 nm. The objective was a 25× 0.95 NA water immersion lens (Leica). Functional images (512 × 256 pixels) were acquired using ScanImage 5 (vidriotechnologies.com) at 7.5 Hz. Approximate laser power at the sample were measured using a slide power meter (Thorlabs) and 3 mw and 20 mW were used for functional imaging. Trains of 1, 2, 5, 10 and 20 field stimuli (1 ms pulse width at 50 Hz) were applied with a stimulator (NPI ISO-STIM). Stimulation voltage was calibrated to elicit an identifiable response to a single pulse in Rohon-Beard cells without stimulating muscle cells. ROIs were selected manually, and data were analyzed using MATLAB (MathWorks).

### Mouse V1 imaging

For *in vivo* mouse V1 imaging, the procedure was done as previously reported^24^. Briefly, AAV injection was used for expression of K-GECO1 in Mouse V1 neurons. After the virus injection, cranial window was implanted. The animal was then placed under a microscope at 37°C and anesthetized during imaging. A custom-built 2-photon microscope was used for imaging with a pulse laser as light source and a 16× 0.8 NA water immersion lens as objective. Laser power was 100-150 mW at the front aperture of the objective lens. Moving grating stimulus trial consisted of a blank period followed by a drifting sinusoidal grating with eight drifting directions with 45° separation. The gratings were presented with an LCD screen placed in front of the center of the right eye of the mouse. For fixed tissue analysis, mice were anesthetized and transcardially perfused. The brains were then removed and post-fixed. Sections of the brains were coverslipped and imaged using confocal microscopy (LSM 710, Zeiss).

### Statistical analysis

All data are expressed as means ± s.d. Sample sizes (n) are listed for each experiment. For V1 functional imaging, ANOVA test (p=0.01) was used to identify responsive cells for each of the grating stimuli.

## Acknowledgements

We thank the University of Alberta Molecular Biology Services Unit for technical support and Christopher W. Cairo for providing access to instrumentation. YS was supported by an Alberta Innovate Scholarship. ASA was supported by a Vanier Canada Graduate Scholarship and an Alberta Innovates Health Solutions (AIHS) Studentship. BR was supported by NSERC (RGPIN-2014-06484). KB was supported by University Hospital Foundation. Work in the lab of REC is supported by grants from CIHR (MOP-123514), NSERC (RGPIN 288338-2010), Brain Canada, and NIH (U01 NS094246 and UO1 NS090565).

## Author contributions

YS LW MDW performed rational design and directed evolution of K-GECO variants. YS YQ RP performed *in vitro* one-photon characterization. RSM RP MD performed *in vitro* two-photon characterization. ERS crystallized protein and solved the structure. YC performed imaging in HeLa cells. HD ASA NH VR performed imaging in dissociated neurons. YFC performed imaging in iPSC-CMs. BR VR performed imaging of organotypic slices. JS MK performed *in vivo* imaging in zebrafishes. HD performed *in vivo* imaging in mice. REC ERS DSK KB TEH supervised the research. YS REC wrote the manuscript.

## Materials & Correspondence

Correspondence and requests for materials should be addressed to R.E.C.

## Supplementary Material

**Figure S1.**
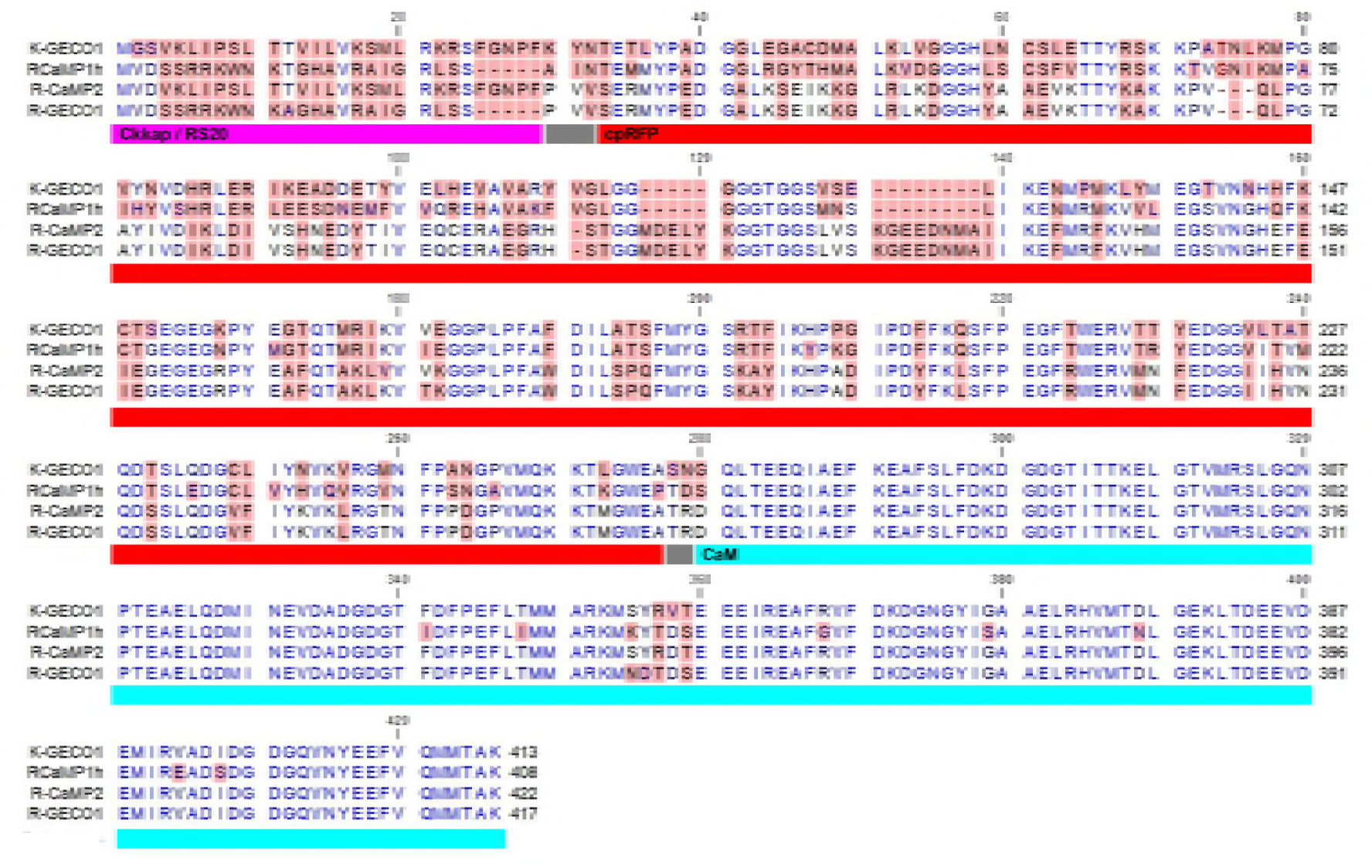
Protein sequence alignment of K-GECO1, R-CaMP2, R-GECO1, and RCaMP1h. Reserved residues are colored in blue. Different residues are highlighted in red. Structural information is indicated with colored bars below the aligned sequences.

**Figure S2.**
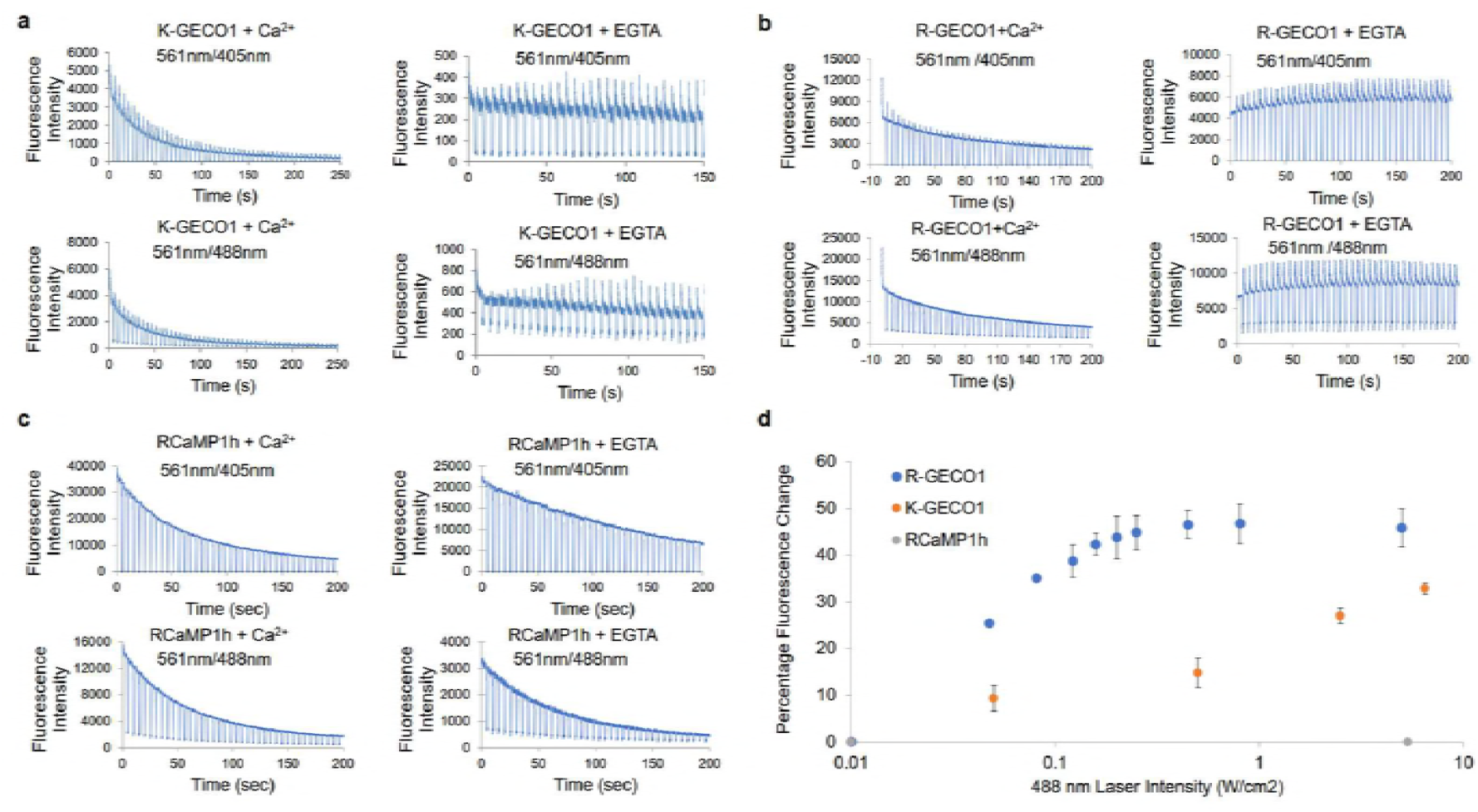
*In vitro* photoactivation characterization of K-GECO1, R-GECO1, and RCaMP1h. **(a)** Representative K-GECO1 fluorescence response to switching between 4 s of illumination with a 561 nm (6.13 W/cm^2^) laser and 1 s with a 405 nm (1.76 W/cm^2^) or 488 nm (6.13 W/cm^2^) laser in the presence and absence (EGTA buffer) of Ca^2+^. **(b)** Representative R-GECO1 fluorescence response with switching between 4 s of 561 nm (3.83 W/cm^2^) laser and 1 s of 405 nm (0.08 W/cm^2^) or 488 nm (3.83 W/cm^2^) laser in both Ca^2+^ buffer and Ca^2+^-free buffer. **(c)** Representative RCaMP1h fluorescence response with switching between 4 s of 561 nm (3.83 W/cm^2^) laser and 1 s of 405 nm (0.08 W/cm^2^) or 488 nm (3.83 W/cm^2^) laser in both Ca^2+^ buffer and Ca^2+^-free buffer. **(d)** Percentage fluorescence change of K-GECO1, R-GECO1, and RCaMP1h in Ca^2+^-free buffer after applying 1 s of 488 nm laser with various intensity when illuminated with 561 nm laser.

**Figure S3.**
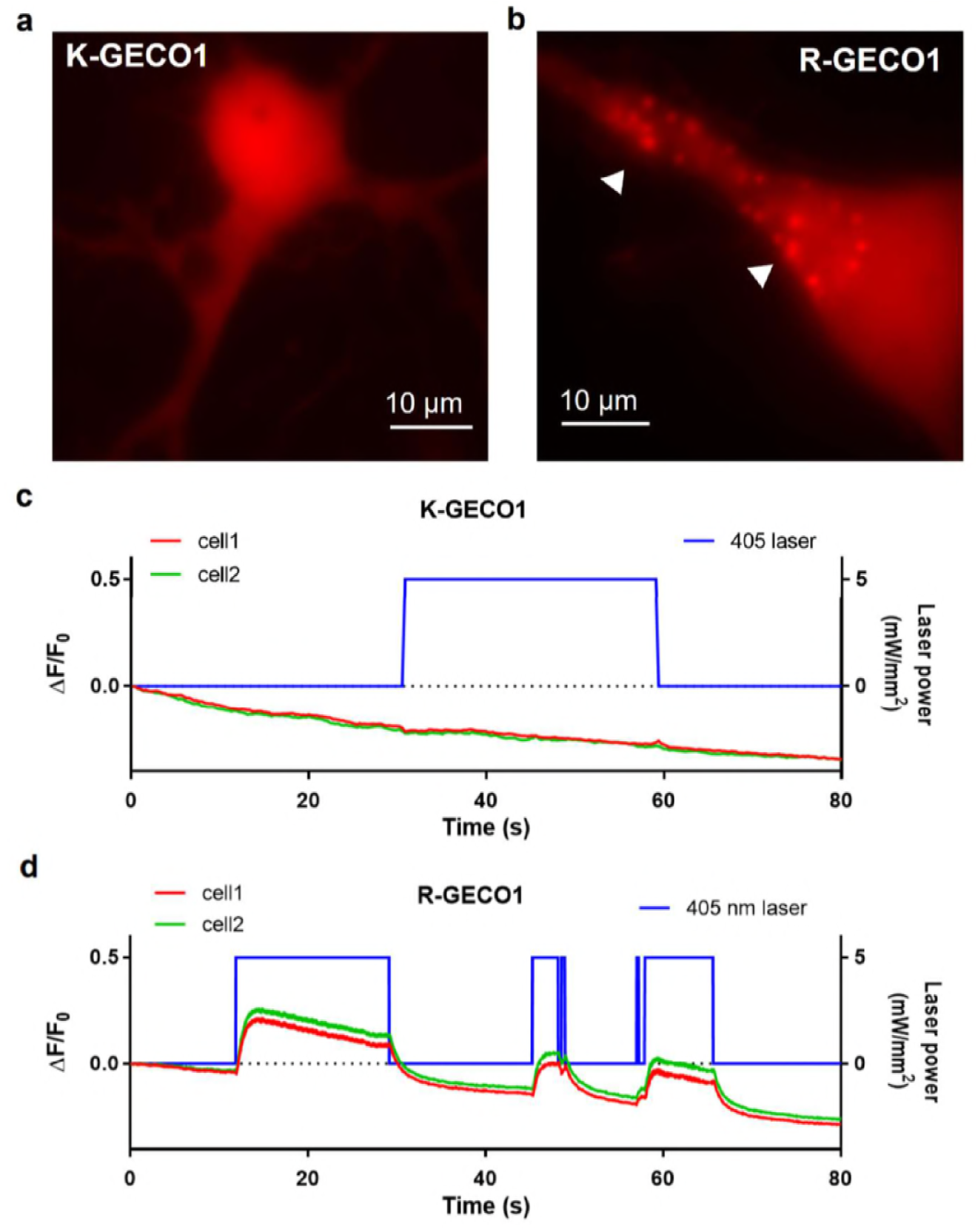
Fluorescence localization and photoactivation of K-GECO1 and R-GECO1 in cultured neurons. (**a**) Representative fluorescence image of K-GECO1 transfected cultured hippocampal neuron. (**b**) Representative fluorescence image of R-GECO1 transfected cultured hippocampal neuron, fluorescent puncta structures are indicated by the arrowhead. (**c**) K-GECO1 fluorescence response in neurons when apply 405 nm laser illumination. (**d**) R-GECO1 fluorescence response in neurons when apply 405 nm laser illumination.

**Figure S5.**
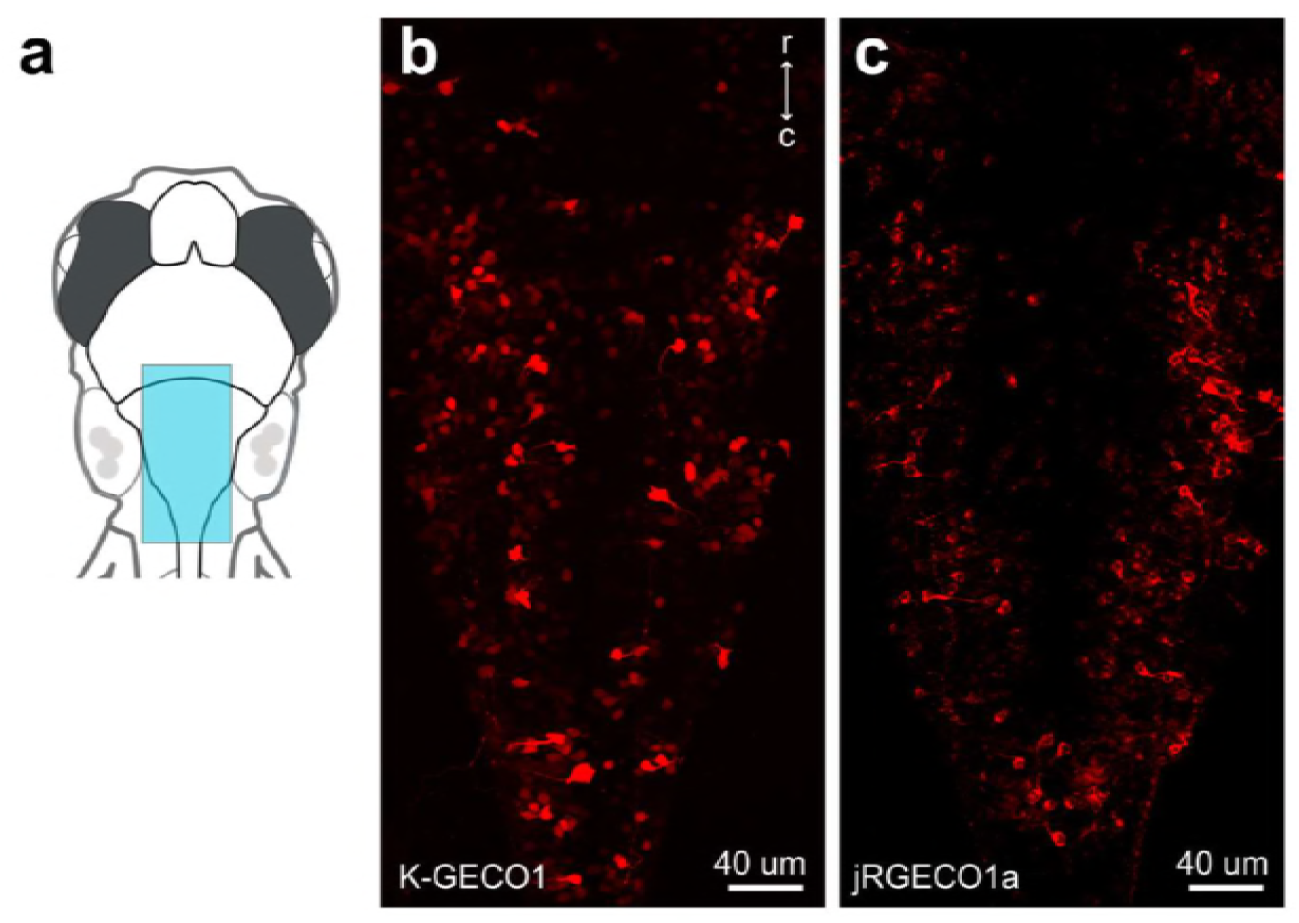
K-GECO1 expression patterns in zebrafish Rohon-Beard (RB) cells. **(a)** Schematic view of the image window. **(b)** Representative images K-GECO1 expression in RB cells. **(c)** Representative images jRGECO1a (with NES) expression in RB cells.

**Figure S5.**
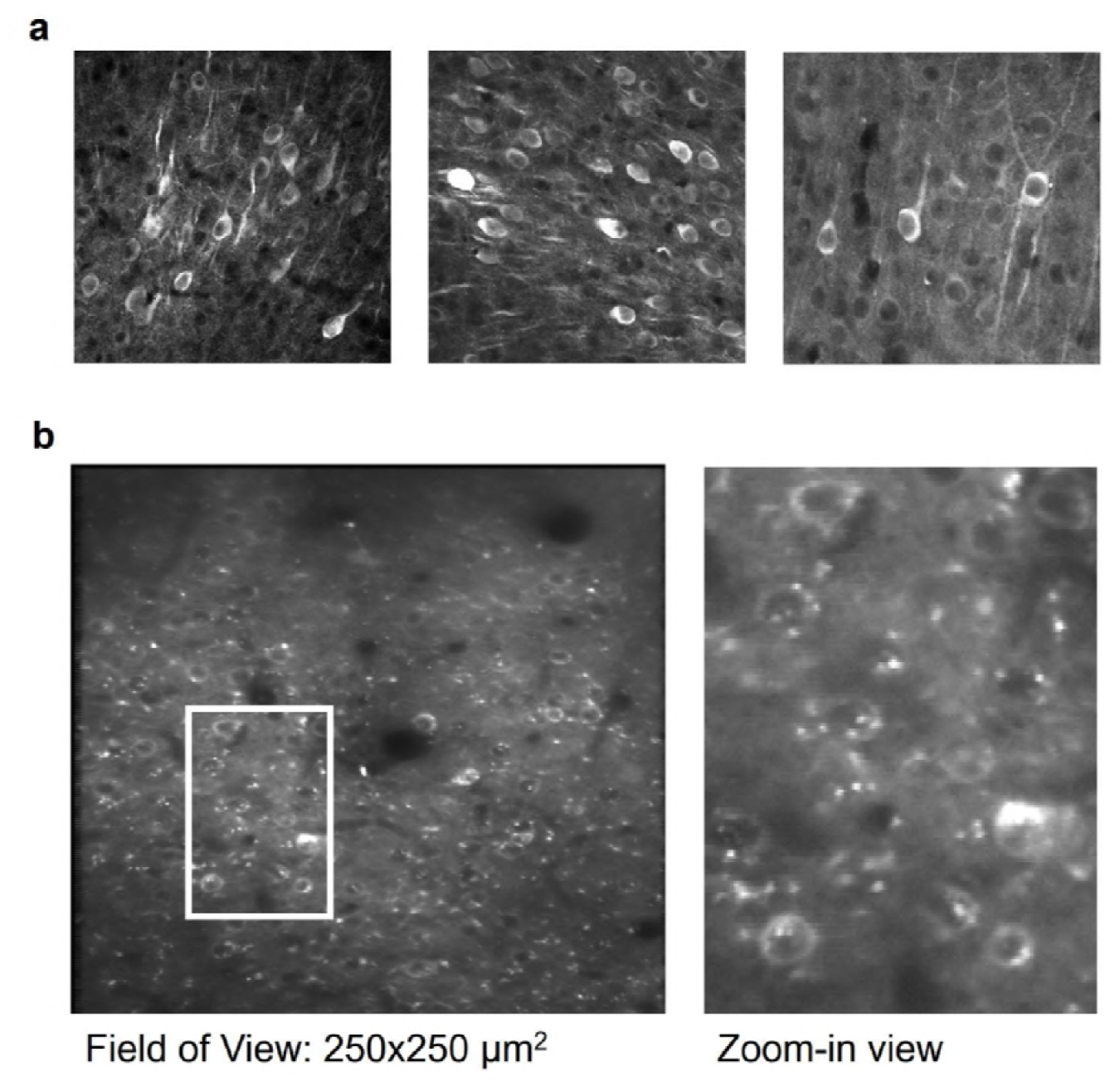
K-GECO1 expression patterns in zebrafish Rohon-Beard (RB) cells. (**a**) Schematic view of the image window. (**b**) Representative images K-GECO1 expression in RB cells. (**c**) Representative images jRGECO1a (with NES) expression in RB cells. (**a**) Representative images K-GECO1 (with NES) expression in fixed tissue section from mouse V1. (**b**) Representative image and zoom-in view of K-GECO1 expression in functional imaging of mouse V1 neurons.

**Supplementary Table S1.** In vitro photophysical characteristics of K-GECO1, R-GECO1, and RCaMP1h (-/+ Ca^2+^).

| | Excitation<br>Maxima<br>(nm) | Emission<br>Maxima<br>(nm) | Intensity<br>fold<br>change | Extinction<br>Coefficient<br>(M<sup>-1</sup>cm<sup>-1</sup>) | Quantum<br>Yield | $K_d$<br>(nM) | Apparent<br>Hill<br>Coefficient | References |
| --- | --- | --- | --- | --- | --- | --- | --- | --- |
| K-GECO1 | 568 (-) | 594 (-) | 12x | 19,000 (-) | 0.12 (-) | 165 | 1.12 | This study |
|  | 565 (+) | 590 (+) |  | 61,000 (+) | 0.45 (+) |  |  |  |
| R-GECO1 | 577 (-) | 600 (-) | 16x | 15,000 (-) | 0.06 (-) | 482 | 2.06 | 18 |
|  | 561 (+) | 589 (+) |  | 51,000 (+) | 0.20 (+) |  |  |  |
| RCaMP1h | 575 (-) | 602 (-) | 12x | 18,700 (-) | 0.14 (-) | 1300 | 2.50 | 19 |
|  | 571 (+) | 594 (+) |  | 65,100 (+) | 0.51 (+) |  |  |  |

**Supplementary Table S2.** X-ray diffraction data collection and model refinement statistics.

|  | **K-GECO**<br>**PDB ID 5UKG** |
| --- | --- |
| **Data collection** |  |
| Space group | P2<sub>1</sub> |
| Unit cell dimensions |  |
| a (Å) | 42.3 |
| b (Å) | 129.2 |
| c (Å) | 78.7 |
| b (deg) | 106.3 |
| Beamline | ALS 8.2.1 |
| Wavelength (Å) | 1.000 |
| Resolution range (Å) | 50 – 2.4 |
| Total reflections | 146,057 |
| Unique reflections | 30,298 |
| Completeness (%)<sup>a</sup> | 94.6 (74.1) |
| I/σ<sup>a</sup> | 5.1 (1.3) |
| R<sub>sym</sub> (%)<sup>a,b</sup> | 20.1 (58.1) |
| CC<sub>1/2</sub> in highest resolution shell | 0.697 |
| **Refinement** |  |
| R<sub>work</sub> / R<sub>free</sub> (%)<sup>c</sup> | 21.0/27.5 |
| Resolution range (Å) | 76 – 2.36 |
| Number of atoms (B factor) |  |
| protein | 6203 (36.9) |
| Ca<sup>2+</sup> | 8 (41.8) |
| water | 64 (31.4) |
| RMSD values |  |
| Bond lengths (Å) | 0.012 |
| Bond angles (degrees) | 1.71 |
| Ramachandran (%) |  |
| Favored/Disallowed | 97.1/0.4 |
| Molprobity |  |
| Clashscore (percentile) | 2.4 (100) |
| Molprobity score (percentile) | 1.44 (99) |

<sup>a</sup>The number in parentheses is for the highest resolution shell.
$R_{\text{sym}} = \sum_{hkl} |I^i(hkl) - \langle I(hkl) \rangle| / \sum_{hkl} \langle I(hkl) \rangle$ , where $I^i(hkl)$ is the $i^{\text{th}}$ measured diffraction intensity and $\langle I(hkl) \rangle$ is the mean of the intensity for the miller index $(hkl)$ .
$R_{\text{work}} = \sum_{hkl} ||F_o(hkl)| - |F_c(hkl)|| / \sum_{hkl} |F_o(hkl)|$ . $R_{\text{free}} = R_{\text{work}}$ for 5% of reflections not included in refinement.

## Reference

1. Campbell, R. E., Tour, O., Palmer, A. E., Steinbach, P. A., Baird, G. S., Zacharias, D. A. & Tsien, R. Y. A monomeric red fluorescent protein. Proc. Natl. Acad. Sci. U. S. A. 99, 7877–7882 (2002).

2. Shaner, N. C., Campbell, R. E., Steinbach, P. A., Giepmans, B. N. G., Palmer, A. E. & Tsien, R. Y. Improved monomeric red, orange and yellow fluorescent proteins derived from Discosoma sp. red fluorescent protein. Nat. Biotechnol. 22, 1567–1572 (2004).

3. Shaner, N. C., Lin, M. Z., McKeown, M. R., Steinbach, P. A., Hazelwood, K. L., Davidson, M. W. & Tsien, R. Y. Improving the photostability of bright monomeric orange and red fluorescent proteins. Nat. Methods 5, 545–551 (2008).

4. Shen, Y., Chen, Y., Wu, J., Shaner, N. C. & Campbell, R. E. Engineering of mCherry variants with long Stokes shift, red-shifted fluorescence, and low cytotoxicity. PLoS One 12, e0171257 (2017).

5. Merzlyak, E. M., Goedhart, J., Shcherbo, D., Bulina, M. E., Shcheglov, A. S., Fradkov, A. F., Gaintzeva, A., Lukyanov, K. A., Lukyanov, S., Gadella, T. W. J. & Chudakov, D. M. Bright monomeric red fluorescent protein with an extended fluorescence lifetime. Nat. Methods 4, 555–557 (2007).

6. Wiedenmann, J., Schenk, A., Röcker, C., Girod, A., Spindler, K.-D. & Nienhaus, G. U. A far-red fluorescent protein with fast maturation and reduced oligomerization tendency from Entacmaea quadricolor (Anthozoa, Actinaria). Proc. Natl. Acad. Sci. U. S. A. 99, 11646–11651 (2002).

7. Shcherbo, D., Merzlyak, E. M., Chepurnykh, T. V., Fradkov, A. F., Ermakova, G. V., Solovieva, E. A., Lukyanov, K. A., Bogdanova, E. A., Zaraisky, A. G., Lukyanov, S. & Chudakov, D. M. Bright far-red fluorescent protein for whole-body imaging. Nat. Methods 4, 741–746 (2007).

8. Shcherbo, D., Murphy, C. S., Ermakova, G. V., Solovieva, E. A., Chepurnykh, T. V., Shcheglov, A. S., Verkhusha, V. V., Pletnev, V. Z., Hazelwood, K. L., Roche, P. M., Lukyanov, S., Zaraisky, A. G., Davidson, M. W. & Chudakov, D. M. Farred fluorescent tags for protein imaging in living tissues. Biochem. J 418, 567–574 (2009).

9. Shemiakina, I. I., Ermakova, G. V., Cranfill, P. J., Baird, M. A., Evans, R. A., Souslova, E. A., Staroverov, D. B., Gorokhovatsky, A. Y., Putintseva, E. V., Gorodnicheva, T. V., Chepurnykh, T. V., Strukova, L., Lukyanov, S., Zaraisky, A. G., Davidson, M. W., Chudakov, D. M. & Shcherbo, D. A monomeric red fluorescent protein with low cytotoxicity. Nat. Commun. 3, 1204 (2012).

10. Kredel, S., Oswald, F., Nienhaus, K., Deuschle, K., Röcker, C., Wolff, M., Heilker, R., Nienhaus, G. U. & Wiedenmann, J. mRuby, a bright monomeric red fluorescent protein for labeling of subcellular structures. PLoS One 4, e4391 (2009).

11. Lam, A. J., St-Pierre, F., Gong, Y., Marshall, J. D., Cranfill, P. J., Baird, M. A., McKeown, M. R., Wiedenmann, J., Davidson, M. W., Schnitzer, M. J., Tsien, R. Y. & Lin, M. Z. Improving FRET dynamic range with bright green and red fluorescent proteins. Nat. Methods 9, 1005–1012 (2012).

12. Bajar, B. T., Wang, E. S., Lam, A. J., Kim, B. B., Jacobs, C. L., Howe, E. S., Davidson, M. W., Lin, M. Z. & Chu, J. Improving brightness and photostability of green and red fluorescent proteins for live cell imaging and FRET reporting. Sci. Rep. 6, 20889 (2016).

13. Shen, Y., Lai, T. & Campbell, R. E. Red fluorescent proteins (RFPs) and RFP-based biosensors for neuronal imaging applications. Neurophotonics 2, 031203 (2015).

14. Kim, T. H., Zhang, Y., Lecoq, J., Jung, J. C., Li, J., Zeng, H., Niell, C. M. & Schnitzer, M. J. Long-Term Optical Access to an Estimated One Million Neurons in the Live Mouse Cortex. Cell Rep. 17, 3385–3394 (2016).

15. Tian, L., Hires, S. A., Mao, T., Huber, D., Chiappe, M. E., Chalasani, S. H., Petreanu, L., Akerboom, J., McKinney, S. A., Schreiter, E. R., Bargmann, C. I., Jayaraman, V., Svoboda, K. & Looger, L. L. Imaging neural activity in worms, flies and mice with improved GCaMP calcium indicators. Nat. Methods 6, 875–881 (2009).

16. Akerboom, J., Chen, T.-W., Wardill, T. J., Tian, L., Marvin, J. S., Mutlu, S., Calderón, N. C., Esposti, F., Borghuis, B. G., Sun, X. R., Gordus, A., Orger, M. B., Portugues, R., Engert, F., Macklin, J. J., Filosa, A., Aggarwal, A., Kerr, R. A., Takagi, R., Kracun, S., Shigetomi, E., Khakh, B. S., Baier, H., Lagnado, L., Wang, S. S.-H., Bargmann, C. I., Kimmel, B. E., Jayaraman, V., Svoboda, K., Kim, D. S., Schreiter, E. R. & Looger, L. L. Optimization of a GCaMP calcium indicator for neural activity imaging. J. Neurosci. 32, 13819–13840 (2012).

17. Chen, T.-W., Wardill, T. J., Sun, Y., Pulver, S. R., Renninger, S. L., Baohan, A., Schreiter, E. R., Kerr, R. A., Orger, M. B., Jayaraman, V., Looger, L. L., Svoboda, K. & Kim, D. S. Ultrasensitive fluorescent proteins for imaging neuronal activity. Nature 499, 295–300 (2013).

18. Zhao, Y., Araki, S., Wu, J., Teramoto, T., Chang, Y.-F., Nakano, M., Abdelfattah, A. S., Fujiwara, M., Ishihara, T., Nagai, T. & Campbell, R. E. An expanded palette of genetically encoded Ca2+ indicators. Science 333, 1888–1891 (2011).

19. Akerboom, J., Carreras Calderón, N., Tian, L., Wabnig, S., Prigge, M., Tolö, J., Gordus, A., Orger, M. B., Severi, K. E., Macklin, J. J., Patel, R., Pulver, S. R., Wardill, T. J., Fischer, E., Schüler, C., Chen, T.-W., Sarkisyan, K. S., Marvin, J. S., Bargmann, C. I., Kim, D. S., Kügler, S., Lagnado, L., Hegemann, P., Gottschalk, A., Schreiter, E. R. & Looger, L. L. Genetically encoded calcium indicators for multi-color neural activity imaging and combination with optogenetics. Front. Mol. Neurosci. 6, 2 (2013).

20. Ohkura, M., Sasaki, T., Kobayashi, C., Ikegaya, Y. & Nakai, J. An improved genetically encoded red fluorescent Ca2+ indicator for detecting optically evoked action potentials. PLoS One 7, e39933 (2012).

21. Wu, J., Liu, L., Matsuda, T., Zhao, Y., Rebane, A., Drobizhev, M., Chang, Y.-F., Araki, S., Arai, Y., March, K., Hughes, T. E., Sagou, K., Miyata, T., Nagai, T., Li, W.-H. & Campbell, R. E. Improved orange and red Ca2+ indicators and photophysical considerations for optogenetic applications. ACS Chem. Neurosci. 4, 963–972 (2013).

22. Wu, J., Abdelfattah, A. S., Miraucourt, L. S., Kutsarova, E., Ruangkittisakul, A., Zhou, H., Ballanyi, K., Wicks, G., Drobizhev, M., Rebane, A., Ruthazer, E. S. & Campbell, R. E. A long Stokes shift red fluorescent Ca(2+) indicator protein for two-photon and ratiometric imaging. Nat. Commun. 5, 5262 (2014).

23. Inoue, M., Takeuchi, A., Horigane, S.-I., Ohkura, M., Gengyo-Ando, K., Fujii, H., Kamijo, S., Takemoto-Kimura, S., Kano, M., Nakai, J., Kitamura, K. & Bito, H. Rational design of a high-affinity, fast, red calcium indicator R-CaMP2. Nat. Methods 12, 64–70 (2014).

24. Dana, H., Mohar, B., Sun, Y., Narayan, S., Gordus, A., Hasseman, J. P., Tsegaye, G., Holt, G. T., Hu, A., Walpita, D., Patel, R., Macklin, J. J., Bargmann, C. I., Ahrens, M. B., Schreiter, E. R., Jayaraman, V., Looger, L. L., Svoboda, K. & Kim, D. S. Sensitive red protein calcium indicators for imaging neural activity. Elife 5, e12727 (2016).

25. Shen, Y., Wiens, M. D. & Campbell, R. E. A photochromic and thermochromic fluorescent protein. RSC Adv. 4, 56762–56765 (2014).

26. Shen, Y., Rosendale, M., Campbell, R. E. & Perrais, D. pHuji, a pH-sensitive red fluorescent protein for imaging of exo- and endocytosis. J. Cell Biol. 207, 419–432 (2014).

27. Yamada, Y. & Mikoshiba, K. Quantitative comparison of novel GCaMP-type genetically encoded Ca2+ indicators in mammalian neurons. Front. Cell. Neurosci. 6, 1–9 (2012).

28. Han, L., Zhao, Y., Zhang, X., Peng, J., Xu, P., Huan, S. & Zhang, M. RFP tags for labeling secretory pathway proteins. Biochem. Biophys. Res. Commun. 447, 508–512 (2014).

29. Abdelfattah, A. S., Farhi, S. L., Zhao, Y., Brinks, D., Zou, P., Ruangkittisakul, A., Platisa, J., Pieribone, V. A., Ballanyi, K., Cohen, A. E. & Campbell, R. E. A Bright and Fast Red Fluorescent Protein Voltage Indicator That Reports Neuronal Activity in Organotypic Brain Slices. J. Neurosci. 36, 2458–2472 (2016).

30. Cai, D., Cohen, K. B., Luo, T., Lichtman, J. W. & Sanes, J. R. Improved tools for the Brainbow toolbox. Nat. Methods 10, 540–547 (2013).

31. Pletnev, S., Shcherbo, D., Chudakov, D. M., Pletneva, N., Merzlyak, E. M., Wlodawer, A., Dauter, Z. & Pletnev, V. A crystallographic study of bright far-red fluorescent protein mKate reveals pH-induced cis-trans isomerization of the chromophore. J. Biol. Chem. 283, 28980–28987 (2008).

32. Nakai, J., Ohkura, M. & Imoto, K. A high signal-to-noise Ca2+ probe composed of a single green fluorescent protein. Nat. Biotechnol. 19, 137–141 (2001).

33. Shui, B., Wang, Q., Lee, F., Byrnes, L. J., Chudakov, D. M., Lukyanov, S. A., Sondermann, H. & Kotlikoff, M. I. Circular permutation of red fluorescent proteins. PLoS One 6, e20505 (2011).

34. Truong, K., Sawano, A., Mizuno, H., Hama, H., Tong, K. I., Mal, T. K., Miyawaki, A. & Ikura, M. FRET-based in vivo Ca2+ imaging by a new calmodulin-GFP fusion molecule. Nat. Struct. Biol. 8, 1069–1073 (2001).

35. Mütze, J., Iyer, V., Macklin, J. J., Colonell, J., Karsh, B., Petrášek, Z., Schwille, P., Looger, L. L., Lavis, L. D. & Harris, T. D. Excitation spectra and brightness optimization of two-photon excited probes. Biophys. J. 102, 934–944 (2012).

36. Kurokawa, H., Osawa, M., Kurihara, H., Katayama, N., Tokumitsu, H., Swindells, M. B., Kainosho, M. & Ikura, M. Target-induced conformational adaptation of calmodulin revealed by the crystal structure of a complex with nematode Ca2+/calmodulin-dependent kinase kinase peptide1. J. Mol. Biol. 312, 59–68 (2001).

37. Tidow, H. & Nissen, P. Structural diversity of calmodulin binding to its target sites. FEBS J. 280, 5551–5565 (2013).

38. Palmer, A. E. & Tsien, R. Y. Measuring calcium signaling using genetically targetable fluorescent indicators. Nat. Protoc. 1, 1057–1065 (2006).

39. Chen, Z. P., Levy, A. & Lightman, S. L. Activation of specific ATP receptors induces a rapid increase in intracellular calcium ions in rat hypothalamic neurons. Brain Res. 641, 249–256 (1994).

40. Higashijima, S.-I., Masino, M. A., Mandel, G. & Fetcho, J. R. Imaging neuronal activity during zebrafish behavior with a genetically encoded calcium indicator. J. Neurophysiol. 90, 3986–3997 (2003).

41. Bethge, P., Carta, S., Lorenzo, D. A., Egolf, L., Goniotaki, D., Madisen, L., Voigt, F. F., Chen, J. L., Schneider, B., Ohkura, M., Nakai, J., Zeng, H., Aguzzi, A. & Helmchen, F. An R-CaMP1.07 reporter mouse for cell-type-specific expression of a sensitive red fluorescent calcium indicator. PLoS One 12, e0179460 (2017).

42. Boyden, E. S., Zhang, F., Bamberg, E., Nagel, G. & Deisseroth, K. Millisecond-timescale, genetically targeted optical control of neural activity. Nat. Neurosci. 8, 1263–1268 (2005).

43. Akerboom, J., Rivera, J. D. V., Guilbe, M. M. R., Malavé, E. C. A., Hernandez, H. H., Tian, L., Hires, S. A., Marvin, J. S., Looger, L. L. & Schreiter, E. R. Crystal Structures of the GCaMP Calcium Sensor Reveal the Mechanism of Fluorescence Signal Change and Aid Rational Design. J. Biol. Chem. 284, 6455–6464 (2009).

44. Wardill, T. J., Chen, T.-W., Schreiter, E. R., Hasseman, J. P., Tsegaye, G., Fosque, B. F., Behnam, R., Shields, B. C., Ramirez, M., Kimmel, B. E., Kerr, R. A., Jayaraman, V., Looger, L. L., Svoboda, K. & Kim, D. S. A neuron-based screening platform for optimizing genetically-encoded calcium indicators. PLoS One 8, e77728 (2013).

45. Baird, G. S., Zacharias, D. A. & Tsien, R. Y. Circular permutation and receptor insertion within green fluorescent proteins. Proc. Natl. Acad. Sci. U. S. A. 96, 11241–11246 (1999).

46. Nagai, T., Sawano, A., Park, E. S. & Miyawaki, A. Circularly permuted green fluorescent proteins engineered to sense Ca2+. Proc. Natl. Acad. Sci. U. S. A. 98, 3197–3202 (2001).

47. Zhao, Y., Abdelfattah, A. S., Zhao, Y., Ruangkittisakul, A., Ballanyi, K., Campbell, R. E. & Harrison, D. J. Microfluidic cell sorter-aided directed evolution of a protein-based calcium ion indicator with an inverted fluorescent response. Integr. Biol. 6, 714–725 (2014).

48. Ai, H.-W., Baird, M. A., Shen, Y., Davidson, M. W. & Campbell, R. E. Engineering and characterizing monomeric fluorescent proteins for live-cell imaging applications. Nat. Protoc. 9, 910–928 (2014).

49. Barnett, L. M., Hughes, T. E. & Drobizhev, M. Deciphering the molecular mechanism responsible for GCaMP6m’s Ca2+-dependent change in fluorescence. PLoS One 12, e0170934 (2017).

50. Makarov, N. S., Drobizhev, M. & Rebane, A. Two-photon absorption standards in the 550–1600 nm excitation wavelength range. Opt. Express, OE 16, 4029–4047 (2008).

51. Studier, F. W. Protein production by auto-induction in high density shaking cultures. Protein Expr. Purif. 41, 207–234 (2005).

52. Otwinowski, Z. & Minor, W. Processing of X-ray diffraction data collected in oscillation mode. Methods Enzymol. 276, 307–326 (1997).

53. McCoy, A. J., Grosse-Kunstleve, R. W., Adams, P. D., Winn, M. D., Storoni, L. C. & Read, R. J. Phaser crystallographic software. J. Appl. Crystallogr. 40, 658–674 (2007).

54. Emsley, P., Lohkamp, B., Scott, W. G. & Cowtan, K. Features and development of Coot. Acta Crystallogr. D Biol. Crystallogr. 66, 486–501 (2010).

55. Winn, M. D., Ballard, C. C., Cowtan, K. D., Dodson, E. J., Emsley, P., Evans, P. R., Keegan, R. M., Krissinel, E. B., Leslie, A. G. W., McCoy, A., McNicholas, S. J., Murshudov, G. N., Pannu, N. S., Potterton, E. A., Powell, H. R., Read, R. J., Vagin, A. & Wilson, K. S. Overview of the CCP4 suite and current developments. Acta Crystallogr. D Biol. Crystallogr. 67, 235–242 (2011).

56. Schneider, C. A., Rasband, W. S. & Eliceiri, K. W. NIH Image to ImageJ: 25 years of image analysis. Nat. Methods 9, 671–675 (2012).

57. Ruangkittisakul, A., Schwarzacher, S. W., Secchia, L., Poon, B. Y., Ma, Y., Funk, G. D. & Ballanyi, K. High sensitivity to neuromodulator-activated signaling pathways at physiological [K+] of confocally imaged respiratory center neurons in on-line-calibrated newborn rat brainstem slices. J. Neurosci. 26, 11870–11880 (2006).

58. Ruangkittisakul, A., Schwarzacher, S. W., Secchia, L., Ma, Y., Bobocea, N., Poon, B. Y., Funk, G. D. & Ballanyi, K. Generation of eupnea and sighs by a spatiochemically organized inspiratory network. J. Neurosci. 28, 2447–2458 (2008).

59. Kimura, Y., Satou, C. & Higashijima, S.-I. V2a and V2b neurons are generated by the final divisions of pair-producing progenitors in the zebrafish spinal cord. Development 135, 3001–3005 (2008).

60. Ding, J., Luo, A. F., Hu, L., Wang, D. & Shao, F. Structural basis of the ultrasensitive calcium indicator GCaMP6. Sci. China Life Sci. 57, 269–274 (2014).

